# An expanded, phenotypically distinct tumor cell subset with low HPV mRNA increases recurrence risk

**DOI:** 10.64898/2026.11.10.762907

**Authors:** Lovely Raghav, Molly R. Patterson, Malay K. Sannigrahi, Pavithra Rajagopalan, Robert M. Brody, Iain M. Morgan, Phyllis A. Gimotty, Jalal B. Jalaly, Ahmed Diab, Sydney M. Schaffer, Noam Auslander, Jean-Nicolas Gallant, Devraj Basu

## Abstract

Human papillomavirus-related (HPV+) cancers harbor a subset of tumor cells expressing certain senescence-related transcripts and reduced levels of HPV mRNAs. We aimed to define this “HPV-low” population’s biologic traits and clinical significance via integrated analysis of recurrence-prone human HPV+ oropharyngeal cancers (OPCs) and patient-derived xenografts (PDXs) of variable therapy sensitivity. Single-cell mRNA sequencing of PDXs and human tumors detected gene expression profiles in HPV-low cells indicating lack of senescence or terminal differentiation and intact tumorigenic capacity. HPV-low cells in cisplatin-refractory PDXs and particular human OPCs differentially upregulated multiple pathways linked to aggressive tumor biology, including RAS signaling, IL-6 signaling, and EMT. Predicted loss of cell-cell interactions among HPV-low cells reflected reduced epithelial identity, further supporting their more aggressive biology. Accordingly, the size of the pre-treatment HPV-low fraction had strong, independent association with recurrence risk. Genetic subclones within OPCs varied widely in content of HPV-low cells, which were enriched in particular lineages. Our findings uncover novel pro-tumor functions of HPV-low cells and support a central role for them in therapy resistance as well as potential utility as a clinical biomarker. Further elucidating mechanisms of cooperativity and plasticity between HPV-low and high states holds promise to reveal targets in treatment-refractory HPV+ malignancies.

## INTRODUCTON

While rates of cervical cancer have gradually declined, human papillomavirus (HPV)-related oropharyngeal carcinomas (OPCs) are rising in incidence. HPV+OPCs have become the most common HPV-related cancer type in the US and UK^1,2^, where HPV+ anal cancers are also increasing, now approaching cervical cancer in incidence^3^. HPV+ OPCs are treated with varying combinations of radiation therapy, cisplatin, and/or surgery, which are curative for the majority but cause serious acute and late toxicities that leave lasting disabilities in survivors^1,2^. In addition, some HPV+ tumors resist such interventions and are predestined to recur. Distinguishing therapy-sensitive and resistant HPV+ cancers at presentation remains challenging and presents a barrier to personalized therapies that maximize cure rates while limiting treatment toxicity.

Varying degrees of intra-tumor heterogeneity in viral gene expression are observed in HPV-driven malignancies and may offer insights into clinically relevant differences in HPV+ cancer biology. The normal HPV life cycle requires spatially regulated shifts in viral oncoprotein levels that modulate keratinocyte self-renewal, proliferation, and differentiation within stratified squamous epithelia^4^. Therefore, analogous variability in levels of the oncogenic drivers encoded by high-risk HPVs is a promising basis to define distinct tumor cell states that cooperatively shape cancer behavior. Maintenance of such heterogeneity among tumor cells in certain HPV+ cancers may confer the phenotypic plasticity needed to progress in hostile microenvironments and resist therapy.

Single cell technologies have yet to be fully leveraged to define the distinct tumor cell states that shape HPV+ cancer progression and therapy responses. A few prior single cell RNA sequencing (scRNAseq) studies characterized heterogeneity in HPV mRNA expression among tumor cells within human OPC specimens^5–8^. One such study analyzed a subset of OPC cells where HPV transcripts were undetectable, showing them to downregulate host mRNAs involved in cell cycle progression and upregulate certain transcripts linked to cellular senescence^7^. A second recent study described a similar HPV+ OPC population by spatial RNAseq, showing it to possess features of the senescence-associated secretory phenotype (SASP)^5^. These cells were enriched in microdomains characterized by tumor hypoxia, increased neutrophils, and CXCL8 chemokine signaling. The functional role of this cell population has not been explored in depth to this date, and the clinical significance of differences in its representation among HPV+ cancers remains unknown.

Relevant prior work is partly limited by scant yields of OPC cells from individual specimens, requiring aggregated analysis of multiple tumors. This approach prevents discrimination of differences among tumors, including features associated with more aggressive OPC behavior. Instead, the contribution of OPC cells with low HPV mRNA to therapy resistance and poor survival was extrapolated from bulk analyses of cell lines and one 28 patient OPC cohort^7^. In addition, low HPV transcript abundance usually results in detection of zero, one, or just a few scRNAseq reads in each OPC cell. The narrow dynamic range of detection has made the dichotomized comparison of OPC cells with zero vs. any HPV reads of unclear biologic significance.

To address limitations of prior studies, we performed scRNAseq on large numbers of tumor cells purified from HPV+ OPC patient-derived xenografts (PDXs) with variable cisplatin sensitivity. For each model, we defined cut points for HPV mRNA levels that maximized host gene expression differences upon dividing OPC cells into “HPV-high” and “HPV-low” groups. Our findings reveal novel differences in host gene expression between the two groups, with HPV-low cells upregulating genes underlying distinct pro-tumor functions. Such features were primarily upregulated in HPV-low cells from those PDXs with innate cisplatin resistance and were also detectable in HPV-low cells from a subset of human OPCs. To assess clinical significance of the HPV-low cell fraction, tumor cells lacking detectable HPV E6/E7 by an RNAscope assay were quantified in a large cohort of HPV+ OPCs that failed standard therapy in comparison to closely matched control tumors that were cured. Our findings reveal a distinct subset of HPV+ OPC cells in which low viral oncogene expression is associated with distinct tumor-promoting functions, relative therapy resistance, and predisposition to recurrence.

## RESULTS

### High cell yields from PDXs with fidelity to the molecular traits and therapy responses of HPV+ OPC

Models were chosen from a panel of HPV+ OPC PDXs that retains key genetic traits of human HPV+ OPCs lost in cancer cell lines^9^. Five PDXs that provided the highest OPC cell yields that are significantly more than prior scRNAseq studies of human HPV+ OPCs (Fig. 1a) were selected for analysis. All five expressed HPV16, and four contained the A1 sub-lineage, which predominates in OPC^10^. Two PDXs (LNT20 and OPTT59) also retained episomal HPV forms^10^, which are seen in ∼50% of human OPCs but lost in HPV+ cancer cell lines^11^. Treating the five PDXs *in vivo* for 12 days (Fig. 1b) produced responses in three, ranging from complete response (CR) in one to stable disease (SD) in two by mRECIST criteria^12^, whereas two models showed progressive disease (PD) during cisplatin therapy (Fig. 1c). No relationship was seen between cisplatin resistance in each model quantified as T/C ratio and bulk levels of HPV E6/E7 or total HPV mRNA (Fig. 1d) defined by prior RNAseq^13^. These results highlight the molecular fidelity and varied therapy responses of the PDX models along with their high cell yields for scRNAseq.

**Figure 1:**
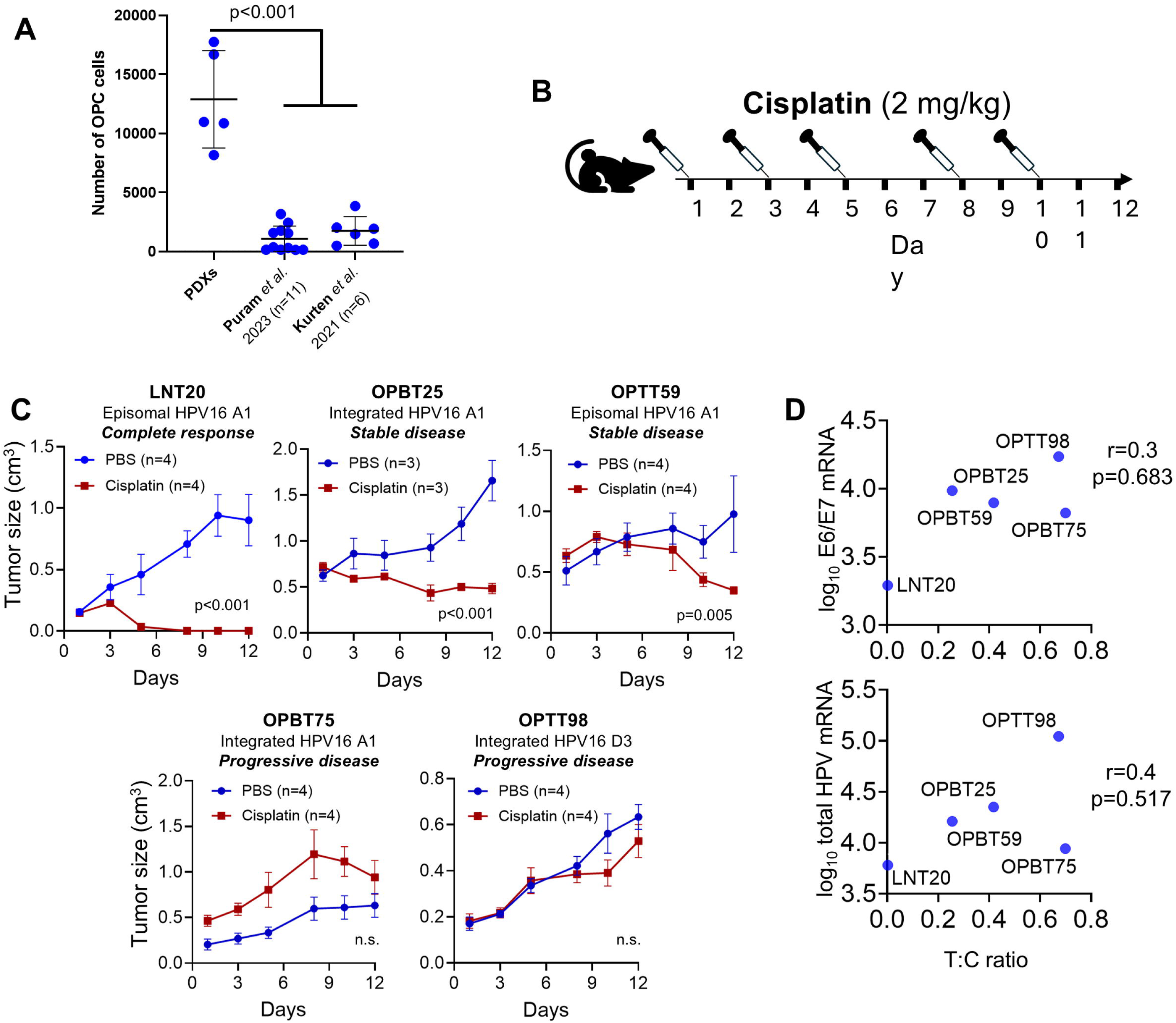
Cell yields, cisplatin responses, and bulk HPV mRNA expression by OPC PDXs. **(A)** HPV+ OPC cell yields from PDXs undergoing scRNAseq in this study relative to HPV+ OPC cells from human samples in two prior studies. P values derived from Dunnett’s multiple comparisons test. **(B)** *In vivo* cisplatin treatment regimen for PDXs grown s.c. in NSG mice. **(C)** Responses of PDXs to cisplatin. HPV16 lineage and genome state are indicated with classification of responses using mRECIST criteria. Error bars represent SEM. P values are from 2-way ANOVA (column effect). **(D)** Absence of correlation between PDX cisplatin responses quantified as T:C ratio and normalized HPV mRNA reads from prior bulk mRNAseq, Spearman correlation coefficients and corresponding P values are shown.

### Optimizing cut points for HPV mRNA levels to dichotomize HPV-high vs. HPV-low cell states

ScRNAseq was performed on PDXs to evaluate relationships between HPV oncoprotein function at the single cell level and therapy resistance. The distribution of HPV mRNA read counts among the OPC cells was used to define optimal expression boundaries for segregating HPV-low cells from HPV-high cells in each PDX. Values across the range of HPV read counts were tested individually as potential cut points to dichotomize the cell population. For this purpose, UMAP embeddings for each group were used to determine “centroid” positions^14^, and cut points providing greatest centroid distance between two groups, as schematized in Fig. 2a, were selected as the boundaries predicted to best differentiate host transcriptional features associated with reduced HPV mRNA. Distributions of both E6/E7 reads and total HPV16 reads were considered, and the latter was selected due to the wider dynamic range provided by total reads in each PDX (Fig, 2b). In two PDXs, ≤1 reads optimally defined the HPV-low state, whereas 0 reads were optimal for the other three (Fig. 2b). This optimization served to increase sensitivity for host gene expression differences associated with HPV mRNA levels, whose dynamic range was constrained by low single cell read counts. Maximizing overall gene expression differences also maximized differences in cancer-related gene expression programs, indicated by the number of number of Hallmark pathways^15^ differentially expressed between groups by GSEA (Fig. 2c). The same pipeline for optimizing HPV expression cut points was applied to the human HPV+ OPC specimens from two prior studies (Fig. 1c), selecting the nine samples that each contained scRNAseq data for at least 1000 OPC cells. For the human samples, 0 vs. 1 HPV reads were consistently the optimal boundary based on centroid distance (Supplementary Fig. 1). Our observations support feasibility of characterizing distinct biologic properties of HPV-low cells in PDXs using HPV expression cut points, despite the paucity of HPV mRNA reads detectable in single OPC cells.

**Figure 2.**
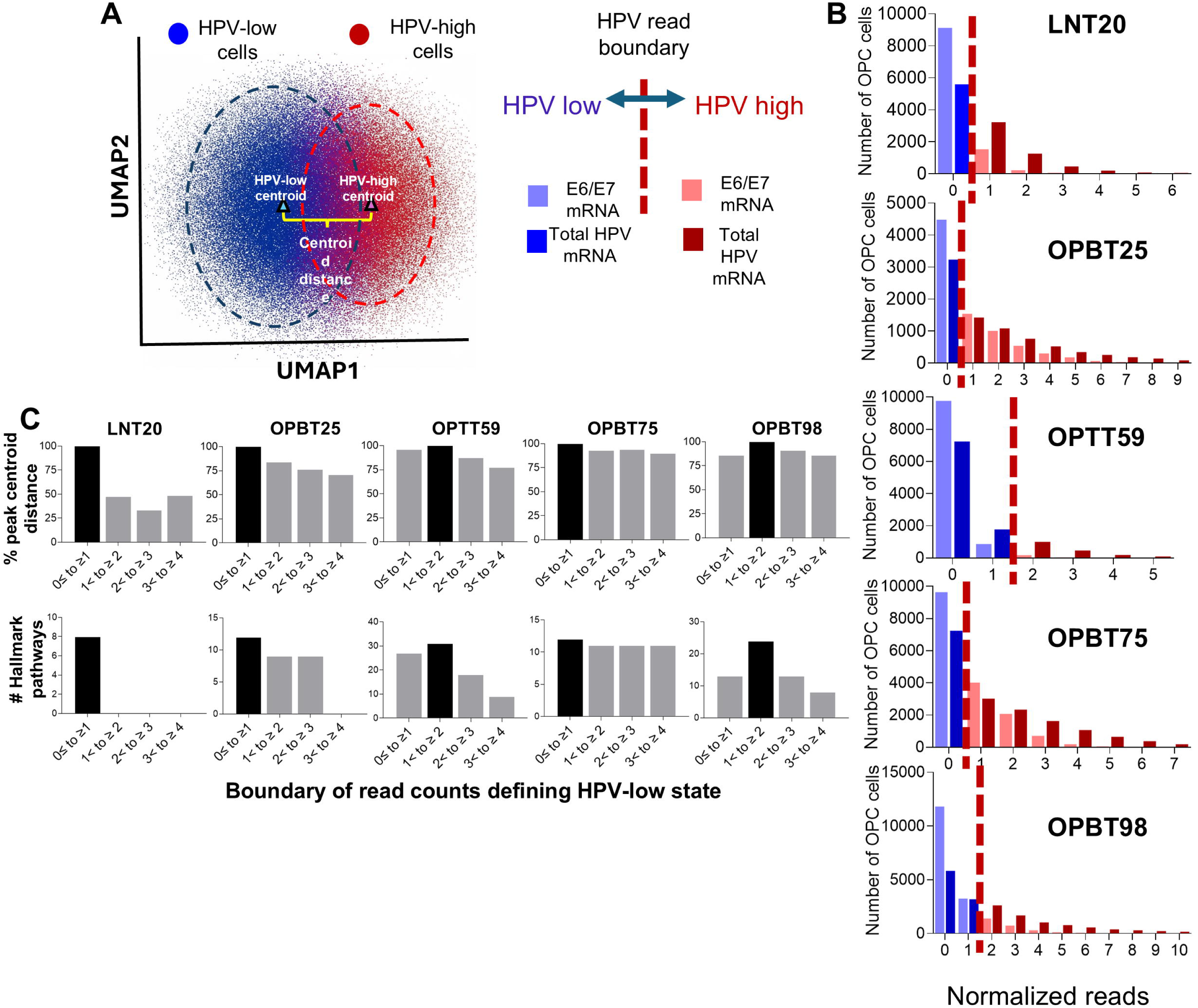
Defining HPV mRNA read cut points that maximize host gene expression differences. **(A)** Schematic illustrating method for measuring centroid distance between OPC cell populations at each HPV mRNA read cut point. **(B)** Distribution of normalized HPV E6/E7 and total mRNA reads across OPC cell populations in PDX models. Dashed lines indicate the read count boundary maximizing centroid distance between HPV low and high subsets for each PDX. **(C)** Concordance between maximized centroid distance (top) and number of differentially expressed Hallmark pathways (bottom). Blue boxes indicate selected read count boundary for each PDX.

### Transcriptional features of HPV-low OPC cells support intact tumor-propagating capacity

To assess relative tumorigenic capacity, HPV-low cells were compared to HPV-high cells for transcriptional features associated with stemness, proliferation, differentiation, and senescence. They were first tested for upregulation of the senescence-related “EpiSen” program^7^ previously observed in HPV+ OPC cells lacking detectable HPV mRNA. HPV-low cells showed marked enrichment of EpiSen program in all five PDXs (Fig. 3a) but did not upregulate the SASP and other senescence-related transcripts previously attributed to HPV-low cells by spatial transcriptomic analysis^5^ (Supplementary Fig. 2a). Although EpiSen contains features of fully developed senescence programs, it is also linked to stem-like cancer cell states with “pre-senescent” gene expression^7^. To distinguish these possibilities, the HPV-low cells were analyzed for features indicative of fully developed senescence. Each PDX OPC cell was classified by the SenePy pipeline^16^, which integrates large-scale scRNAseq data across human tissues and ages to optimize senescence detection. Lower SenPy scores were found in HPV-low cells from all 5 PDXs (Supplementary Fig. 2b). The two most cisplatin-resistant PDXs (OPBT75 and OPTT98) each had three available biologic replicates, facilitating statistical comparison of the percent meeting the full senescence threshold defined by the classifier. This value was unchanged or reduced in HPV-low vs. HPV-high cells of these models (Fig. 3b, left and middle), and a trend toward fewer senescent cells in the HPV-low compartment was evident across all 5 models (Fig. 3b, right). The nine human OPC specimens in aggregate showed markedly fewer senescent cells in their HPV-low fractions (Fig. 3c). Cell states were further mapped using CytoTRACE 2^17^, a deep learning-based predictor of single cell developmental potential that is applicable to cancer cells. CytoTRACE 2 detected abundant oligopotent and unipotent stem-like states in both HPV-high and HPV-low cells of all PDXs as well as rare multipotent cells in one PDX (Supplementary Fig. 2c). A minority fraction of more differentiated cells was present in both subsets and was not increased in the HPV-low fraction of the PDXs when considered in aggregate (Fig. 3d, left) or individually in the two resistant PDXs with biologic replicates (Fig. 3d, right). Likewise, HPV-low cells expressing the epithelial terminal differentiation marker involucrin (*IVL*) mRNA were not increased in the PDXs in aggregate (Fig. 3e, left) and were significantly reduced in one of two resistant PDXs with biologic replicates (Fig. 3e, right). Another terminal differentiation marker *TGM1* showed concordant results (Supplementary Fig. 2d). Together, this analysis supports presence of stem-like states and tumor-propagating capacity in the majority of HPV-low cells along with low vulnerability to mitotic arrest via senescence or terminal differentiation.

**Figure 3.**
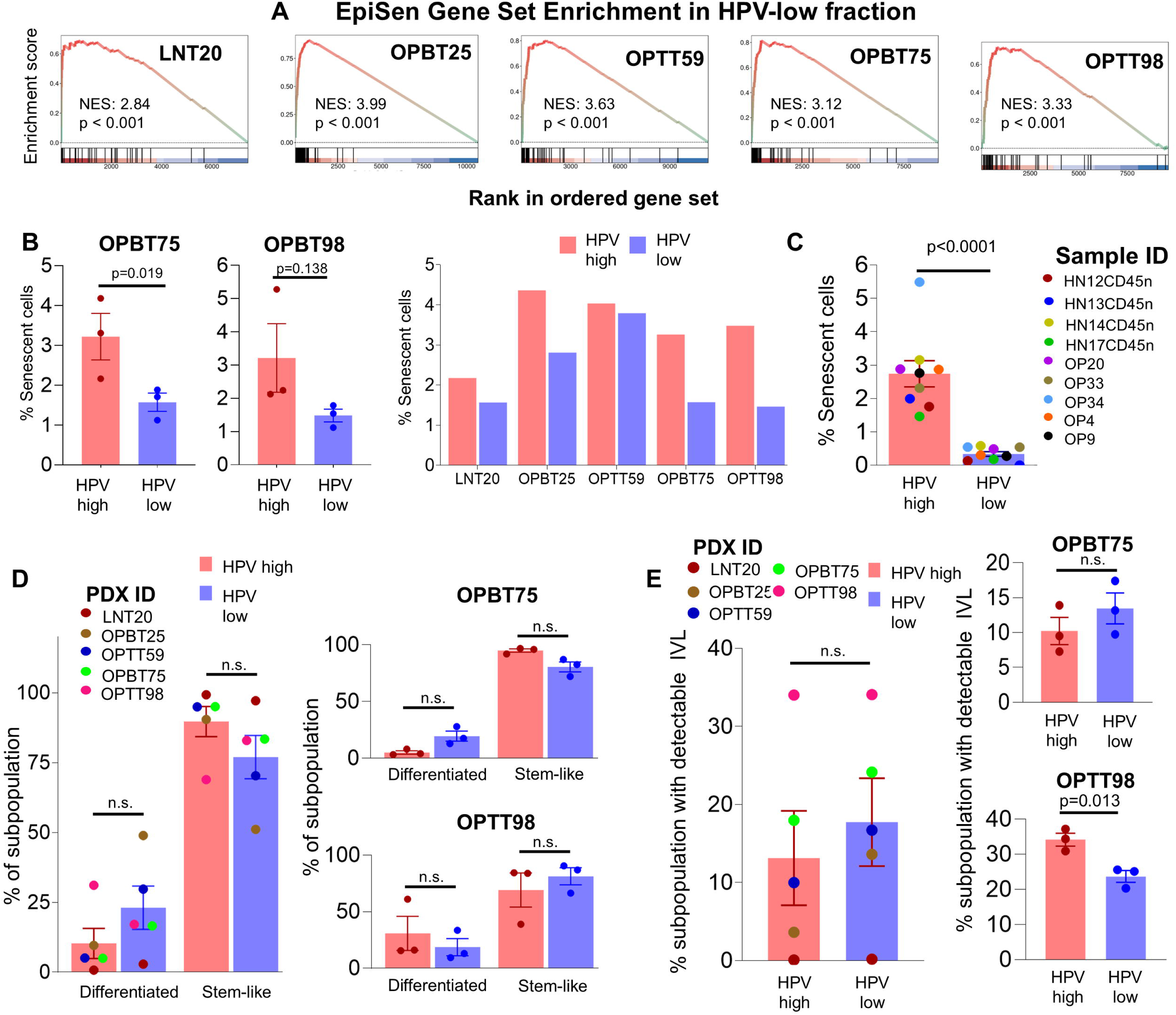
Reduced senescence and intact tumor-propagating capacity in HPV low cells. **(A)** GSEA of the EpiSen program genes in HPV low versus HPV high cells in the 5 PDX models, with p values derived by Benjamini–Hochberg false discovery method. **(B)** Senescent cell content defined by SenPy in HPV high vs. low fractions of OPBT75 and OPBT98, the two PDXs with three biologic replicates (left and middle) compared by paired t-test and across all PDXs, pooling cells from replicates where available (right). **(C)** Senescent fraction sizes in HPV high vs. low cells from primary human OPCs, with p value derived from paired t-test. **(D)** Stem-like vs. differentiated OPC cell fraction sizes mapped by CYTOTRACE2 in HPV-high vs. low subsets, combining all 5 PDX models (left) and using individual replicates of OPBT75 and OPBT98 (right). P values are based on Sidak’s multiple comparisons test. **(E)** Involucrin (IVL) expression in HPV-high vs. low cells across all PDXs (left) and using three replicates of OPBT75 and OPBT98 (right). P values are based on unpaired t tests with Welch’s correction.

### ScRNAseq profiles support a distinct role for HPV-low cells in OPC progression and therapy response

An expanded analysis of cancer-related differences in gene expression between HPV-high and HPV-low populations was performed by GSEA of Hallmark and KEGG pathways, with gene rankings derived from MAST-reported log_2-_fold changes. Pathways differentially expressed between HPV-high and HPV-low states in any PDX are listed in Supplementary Fig. 3a and b, and those significantly downregulated in HPV-low cell vs. HPV-high cells of at least 3 of 5 PDXs are shown in Fig. 4a. The HPV-low fraction in all 5 PDXs had less E2F target gene expression, consistent with less HPV E7 driving cell division through Rb inhibition. Accordingly, the other four recurrently downregulated pathways were involved in cell cycle progression and/or the bioenergetic and biosynthetic demands imposed by rapid proliferation and biomass generation. These pathways upregulated in the HPV-low cells of at least 3 PDXs (Fig. 4b) were also biologically coherent. HPV-low cells in the most cisplatin-sensitive PDX (LNT20) did not upregulate any of these pathways, but the other four upregulated p53 target gene expression, consistent with reduced HPV E6 contributing to higher p53 levels in this population. Upregulated p53 target genes largely explained upregulation of the Hallmark apoptosis gene set in HPV-low cells, suggesting that ongoing apoptosis is not a dominant feature of the HPV-low state. Specifically, removal of p53 target genes from the apoptosis gene list abrogated significance of the latter pathway in 3 of 4 models, despite only 10% overlap between the two Hallmark pathway gene lists (Supplementary Table 1). The other gene sets recurrently upregulated in HPV-low PDX cells included pro-tumorigenic TNFα signaling via NFκB, an oncogenic pathway characteristic of many HPV+ OPCs^18^. Additional upregulated pro-tumorigenic pathways were less characteristic of HPV+ OPC but linked to more aggressive cancer types, specifically KRAS signaling, IL6/JAK/STAT3 signaling, myogenesis, and epithelial mesenchymal transition (EMT). Upregulation of pro-inflammatory pathways was also observed in HPV-low cells and may arise from decreased activity of HPV oncogenes, which can suppress innate and adaptive immunity^19–21^, and/or reflect pro-inflammatory NFκB signaling^22^. The number of upregulated pathways in HPV-low cells tightly correlated across the five PDXs with degree of cisplatin resistance quantified by T/C ratio (Fig. 4c, left), indicating the HPV-low state to be most distinct transcriptionally in the models that progress during cisplatin therapy (OPBT75 and OPTT98). Furthermore, enrichment scores for individual pathways correlated with the T/C ratios for 8 of 17 pathways differentially expressed between HPV-high and HPV-low states in at least 3 PDXs (Fig. 4c, right). Features of HPV-low cells were validated in the human HPV+ OPC panel after filtering for the samples that both contained >1000 OPC cells and upregulated E2F target genes in the HPV-high fraction (n=5). The pathways that were upregulated in HPV-high cells (Fig. 4d) and HPV-low cells (Fig. 4e) in at least 2 of these 5 specimens highly overlapped with those identified in the PDXs (Fig. 4f). Together, these analyses provide evidence of major biologic distinctions between HPV-high and HPV-low compartments in cisplatin-resistant OPCs, with the former linked more directly to tumor growth and the latter potentially serving migration, invasion, and metastasis.

**Figure 4.**
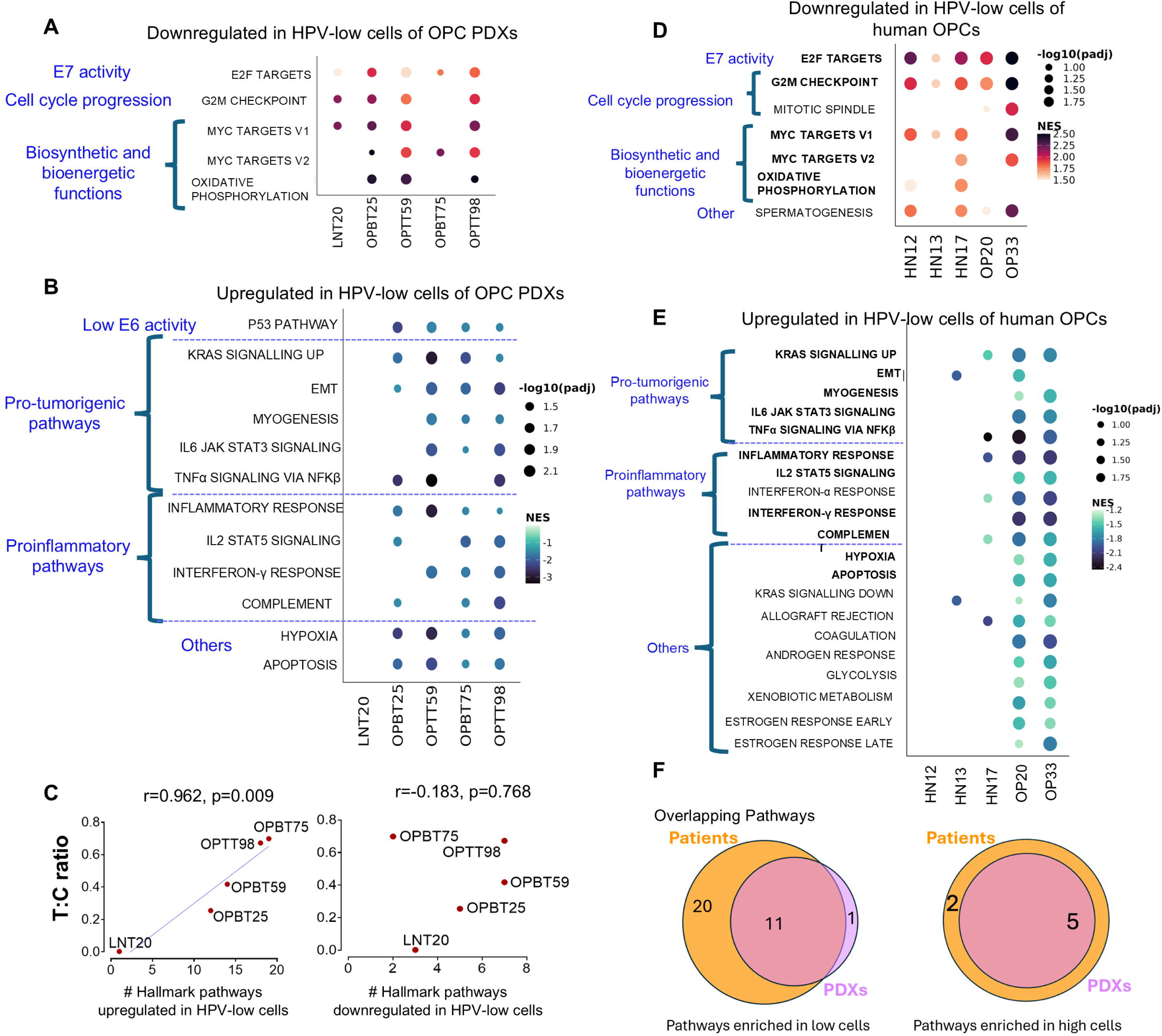
HPV low OPC cells appear less proliferative but upregulate distinct pro-tumor functions. GSEA comparing pseudobulk aggregates of HPV low to HPV high populations was performed using the Hallmark and KEGG pathways. Dot sizes reflect adjusted P values, and color intensity shows normalized enrichment score (NES). Differential pathway expression is designated based on adjusted p<0.05 by the Benjamini–Hochberg method. **(A)** Pathways downregulated or **(B)** upregulated in HPV-low OPC cells in at least 3 of 5 PDX models. **(C)** Correlation analyses of cisplatin T:C ratio with number of pathways enriched in HPV low cells (left) and HPV-high cells (right) of PDXs based on Pearson’s coefficient. **(D)** Pathways downregulated or **(E)** upregulated in HPV-low tumor cells in at least 2 of 5 human HPV+ OPCs. **(F)** Venn diagrams show overlap in pathways characterizing HPV-low cells between PDXs and human tumor samples

### Intra-tumor genetic clones maintain variable HPV expression but differ in HPV-low state abundance

We next evaluated the contribution of clonal heterogeneity within OPCs to representation of the HPV-low cell state by defining tumor cell clones within PDXs using CopyKAT^23^. This tool infers copy number variants in each cell by detecting coordinated up or downregulation of gene expression across broad chromosomal segments and then aggregating individual cell profiles into groups of clones by k-means clustering. This analysis focused on the OPBT75 and OPTT98 PDXs, given that they had the most transcriptionally distinct HPV-low fractions and greatest cisplatin resistance. Six clonal lineages were clustered in OPBT75, and four were clustered in OPTT98 (Fig. 5a). A wide range of HPV gene expression was observed in all clones, suggesting that plasticity in expression is maintained across clonal lineages (Fig. 5b). However, comparing clones as pseudo-bulk entities to each other highlighted different overall HPV expression levels. Upon ordering them from lowest to highest in both PDXs (Fig. 5c), the lowest expression clone (C1) did not differ significantly in HPV mRNA from the second lowest (C2) but diverged from the rest, which were designated as the “C-high” clones in aggregate. The pathways upregulated and downregulated in C1 vs C-high lineages (Fig. 5d) largely recapitulated the functional distinctions attributed to the HPV-low state upfront (Fig. 4a, 4b). These findings indicate HPV expression to vary widely within subclonal populations while also revealing that OPC cells within certain clonal lineages have greater propensity to enter the HPV-low state.

**Figure 5.**
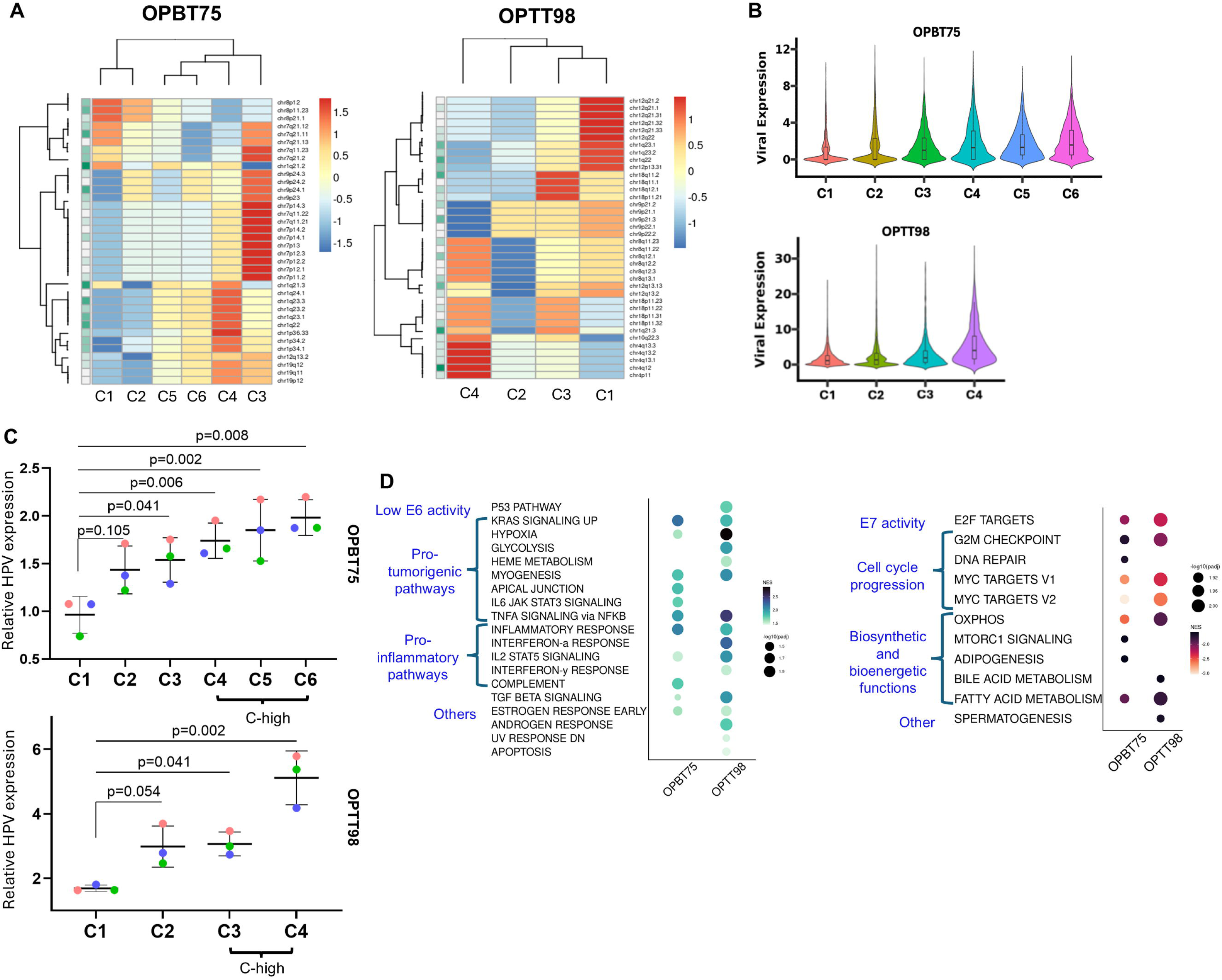
Clonal heterogeneity within HPV+ OPCs contributes to variability in HPV gene expression. **(A)** Non-hierarchical clustering distinguishes clonal lineages in OPBT75 (n=6, labeled C1-C6) and OPTT98 (n=4, labeled C1-4) based on cytobands of copy gains and losses inferred from CopyKAT analysis of scRNAseq. Color gradients represent row-scaled relative copy numbers. **(B)** Distinct distributions of HPV read counts among OPC cells in each clonal lineage within OPBT75 and OPTT98 PDXs. **(C)** Differences in HPV expression among clonal lineages analyzed as pseudobulk entities using three biologic replicates for each PDX. Lineages that differed significantly from the lowest expressing group (C1) are designated as “C-high.” P values were obtained from a one-way ANOVA with post-hoc Dunnett’s multiple comparisons test. **(D)** GSEA showing Hallmark and KEGG pathways significantly downregulated (right) and upregulated (left) in C1 vs. C-high cells. Significance was defined based on adjusted P < 0.05 using the Benjamini–Hochberg method.

### HPV-low and high cell states are jointly maintained during cisplatin responses

The two PDXs (OPBT75 and OPTT98) that progress during cisplatin therapy were further analyzed for relative contributions of HPV-low and HPV-high cells to their resistant phenotypes. To provide insight into the OPC cell state dynamics upon initial cisplatin exposure, PDX-bearing mice were treated as previously (Fig. 1b) but analyzed by scRNAseq earlier (day 6) to isolate cell state dynamics upon initial drug exposure from adaptations to chronic therapy. Treated PDXs showed trends toward more senescent cells using the SenPy classifier (Fig. 6a) but not significantly suggesting limited induction of therapy-induced senescence. However, HPV-low fraction sizes were not altered by therapy in either model (Fig. 6b), and HPV expression was unaltered overall at the pseudo-bulk level (Fig. 6c). Cisplatin-induced changes in OPC cell gene expression were tested in OPBT75 by assigning UMAP clusters to pooled treated and control OPC cells (Fig. 6d, top) and comparing distribution of HPV-low cells among the clusters for the two conditions (Fig. 6d, bottom). No significant changes in cluster sizes (Fig. 6e, top) or distribution of HPV-low states among clusters (Fig. 6e, bottom) were observed in treated vs. control OPBT75 samples, and comparable findings were obtained for OPTT98 (Supplementary Fig. 4). Maintenance of HPV-low and high states in stable balance during initial cisplatin exposure rather than enrichment of one subset suggests cooperativity between them in driving cisplatin resistance.

**Figure 6:**
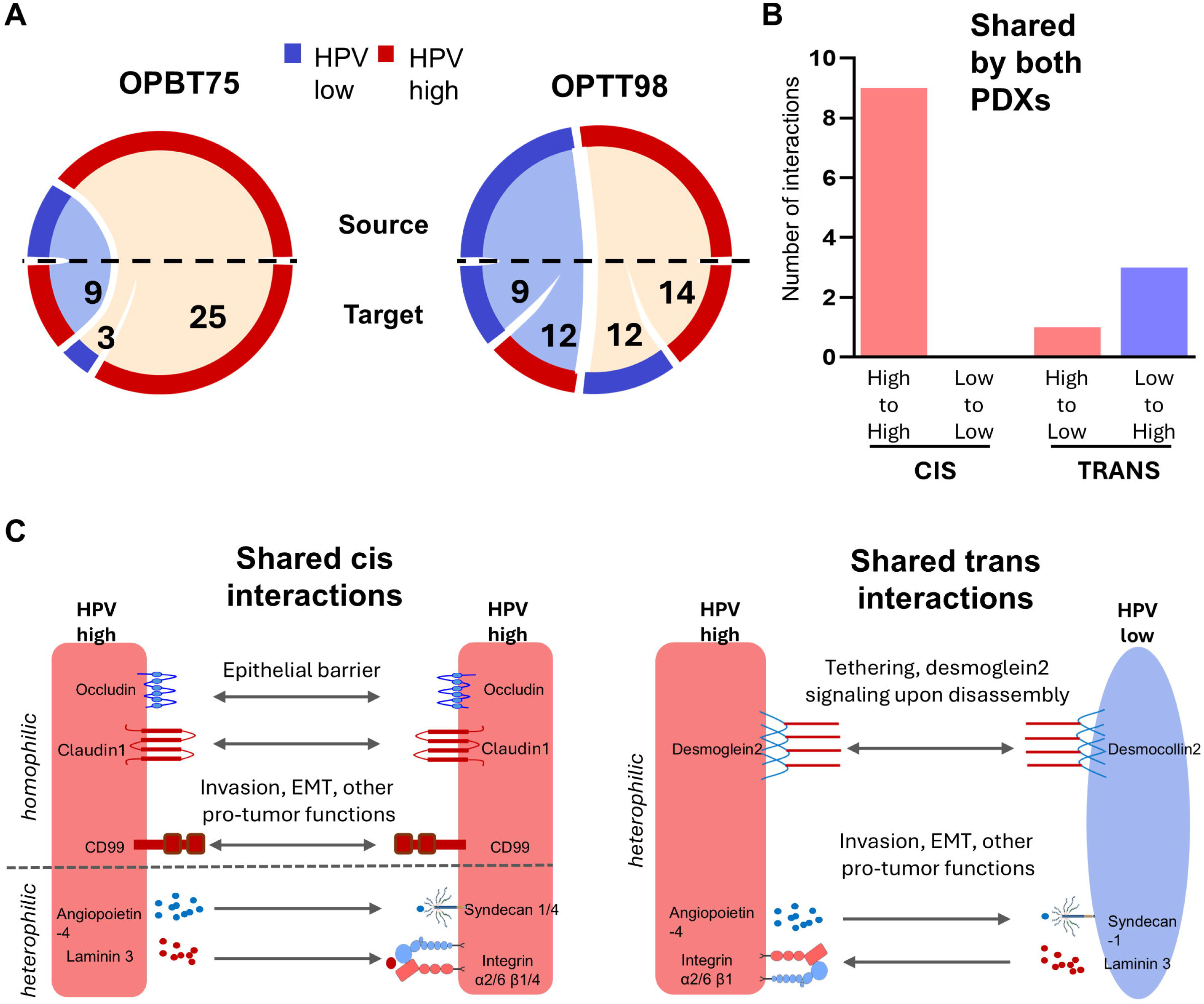
Stability of HPV low and HPV high populations upon treating cisplatin-resistant PDXs. **(A)** Percent senescent cells defined by the SenPy classifier in treated vs. control OPBT75 and OPBT98 PDXs. **(B)** HPV-low fraction sizes in treated vs. control PDXs. **(C)** Pseudo-bulk HPV mRNA reads in treated vs. control PDXs. **(D)** UMAP clustering of treated and control OPBT75 OPC cells (top) overlaid with HPV-low and high cell identities (bottom). **(E)** Proportions of total OPC cells (top) and HPV-low cells (bottom) in each UMAP cluster in control vs. treated OPBT75 PDXs, with lack of significant differences observed using Sidak’s multiple comparisons test. All other comparisons were made using paired t-tests (A-C). Three biological replicates were analyzed for each condition throughout.

### HPV-high cells have greater epithelial identity and potential for complex crosstalk with the HPV-low pool

To infer crosstalk among the HPV-low and high states, we analyzed the extracellular protein-protein interactions predicted among them using CellChat^24^. Significant cis interactions (high/high or low/low) and trans interactions (high/low or low/high) were defined for each cisplatin-resistant PDX (Fig. 7a). These interactions are listed in Supplementary Table 2 and were filtered to those significant in both PDXs (Table 1) and quantified (Fig. 7b). Broad functional classifications for these shared interactions are illustrated in Fig. 7c. The cis interactions were more abundant than trans interactions and only occurred between HPV-high cells. These cis interactions included tight junction components (claudin1-claudin1 and occludin-occludin), supporting greater retention of conventional epithelial functions among HPV-high cells. HPV-high cells also had homophilic cis interactions via CD99, which increases carcinoma cell motility and invasion in some contexts^25^ and thus might participate in HPV-high to low state transition via EMT. Similar effects might be induced by HPV-high cell expression of multiple integrin isoforms that can facilitate migration. HPV-high cells also expressed the secreted factor angiopoietin 4, which can have complex pro-tumor effects through retention on syndecan cell surface proteoglycans^26–28^ expressed by both HPV-high and low cells. Additional crosstalk was predicted through a heterophilic desmosomal protein interaction between desmocollin-2 on HPV-low cells and desmoglein-2 on HPV-high cells. Whereas desmocollins have no signaling domain, desmoglein-2 can signal intracellularly via EGFR/c-Src^29^ upon complex disassembly. Overall, this analysis supports greater epithelial differentiation in the HPV-high state and a role for HPV-high cells in maintaining the HPV-low pool while suggesting complex crosstalk between the subsets contributing to their homeostasis.

**Figure 7.**
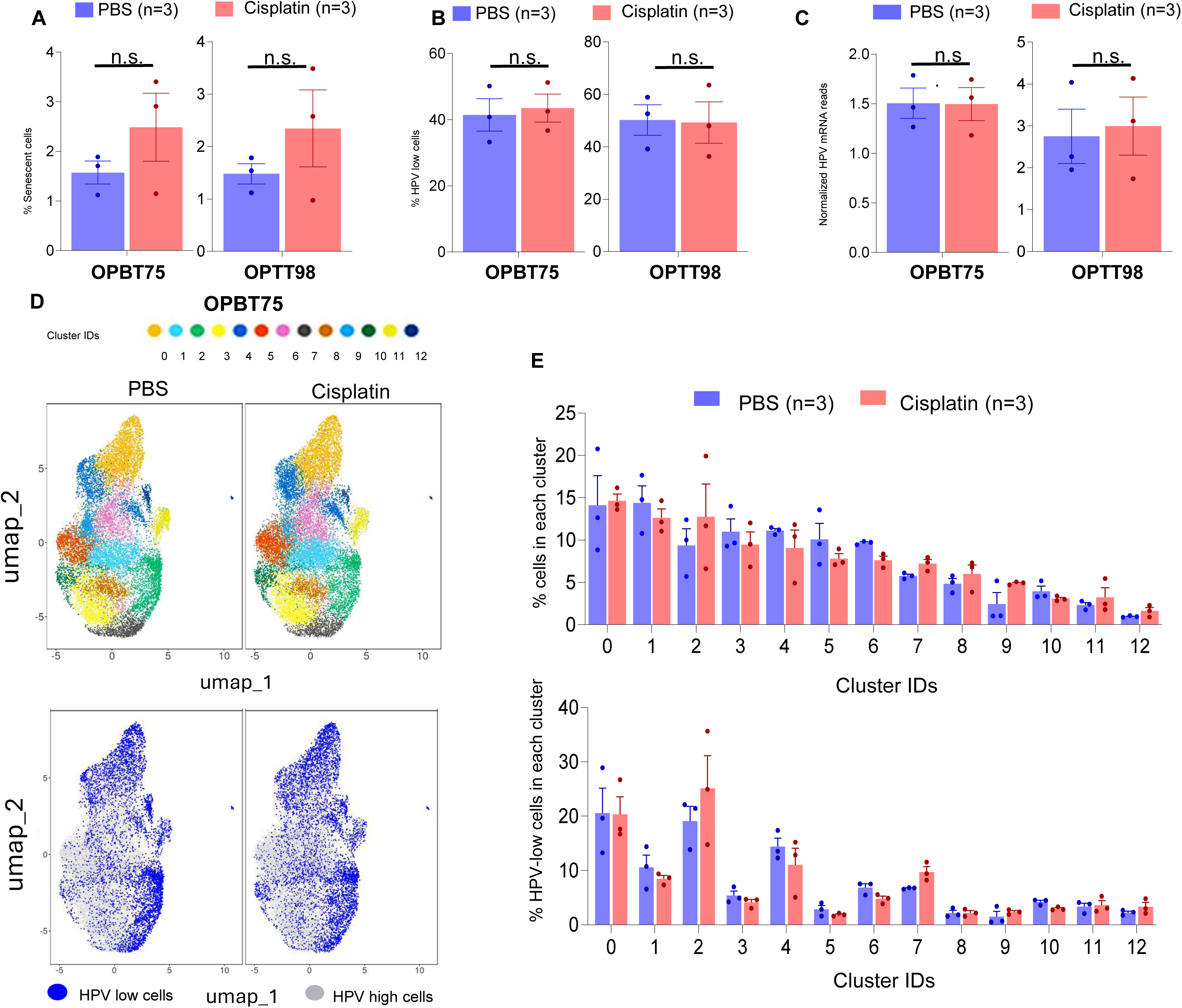
Evidence for cooperation between HPV low and HPV high OPC cell states. **(A)** CellChat analysis of scRNAseq from OPBT75 and OPTT98 (n=3) quantifying potential significant interactions within the HPV-low and HPV-high subsets (cis interactions) and between them (trans interactions), illustrated as chord diagrams **(B)** Quantification of interactions appearing in both OPBT75 and OPTT98. **(C)** Functional groupings of cis and trans interactions appearing in both PDXs.

**Table 1:**
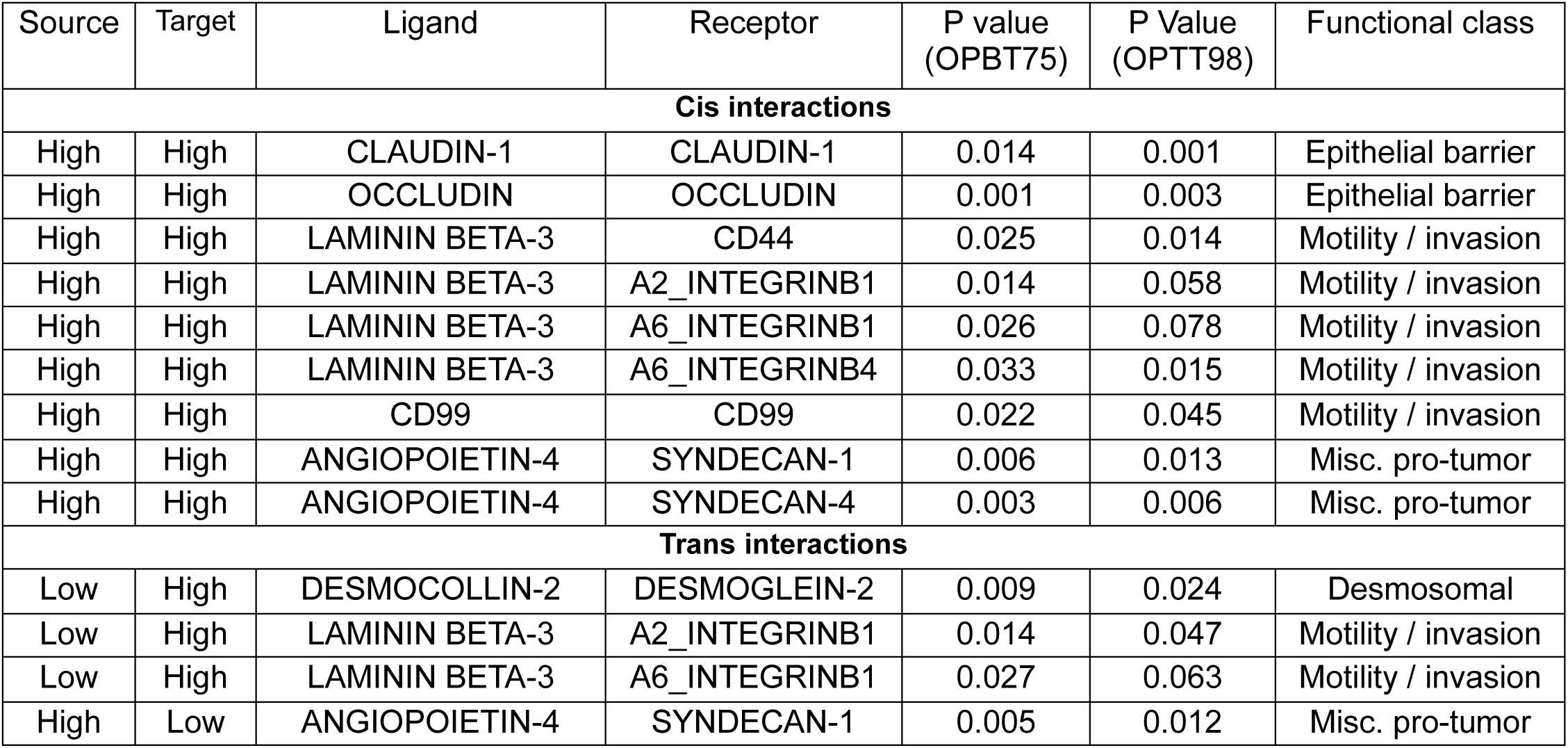
CellChat interactions among HPV-high and low cells shared by OPBT75 and OPTT98. P-values are inferred using triMean method incorporated in CellChat with 10,000 bootstraps.

| Source | Target | Ligand | Receptor | P value<br>(OPBT75) | P Value<br>(OPTT98) | Functional class |
|---|---|---|---|---|---|---|
| **Cis interactions** |  |  |  |  |  |  |
| High | High | CLAUDIN-1 | CLAUDIN-1 | 0.014 | 0.001 | Epithelial barrier |
| High | High | OCCLUDIN | OCCLUDIN | 0.001 | 0.003 | Epithelial barrier |
| High | High | LAMININ BETA-3 | CD44 | 0.025 | 0.014 | Motility / invasion |
| High | High | LAMININ BETA-3 | A2_INTEGRINB1 | 0.014 | 0.058 | Motility / invasion |
| High | High | LAMININ BETA-3 | A6_INTEGRINB1 | 0.026 | 0.078 | Motility / invasion |
| High | High | LAMININ BETA-3 | A6_INTEGRINB4 | 0.033 | 0.015 | Motility / invasion |
| High | High | CD99 | CD99 | 0.022 | 0.045 | Motility / invasion |
| High | High | ANGIOPOIETIN-4 | SYNDECAN-1 | 0.006 | 0.013 | Misc. pro-tumor |
| High | High | ANGIOPOIETIN-4 | SYNDECAN-4 | 0.003 | 0.006 | Misc. pro-tumor |
| **Trans interactions** |  |  |  |  |  |  |
| Low | High | DESMOCOLLIN-2 | DESMOGLEIN-2 | 0.009 | 0.024 | Desmosomal |
| Low | High | LAMININ BETA-3 | A2_INTEGRINB1 | 0.014 | 0.047 | Motility / invasion |
| Low | High | LAMININ BETA-3 | A6_INTEGRINB1 | 0.027 | 0.063 | Motility / invasion |
| High | Low | ANGIOPOIETIN-4 | SYNDECAN-1 | 0.005 | 0.012 | Misc. pro-tumor |

### A larger HPV-low fraction is associated with elevated treatment failure risk in HPV+ OPC patients

HPV-low fraction sizes defined bioinformatically via scRNAseq in the PDXs failed to correlate with cisplatin T/C ratio (not shown). We subsequently tested a more direct approach for defining HPV-low fractions by detecting HPV mRNAs with an E6/E7 RNAScope assay that produces a binary staining result for each OPC cell. This sensitive, clinically validated *in situ* hybridization assay probes for E6/E7 mRNAs encoded by HPV16, 18, 31, and 33^30^ and was used to stain the 5 PDXs harvested at comparable sizes early in their growth curves. Observed correlation in PDXs between the E6/E7 negative (presumptive HPV-low) cell fraction size and cisplatin T/C ratio (Supplementary Fig. 5) encouraged similar evaluation of patients with divergent treatment outcomes. For this purpose, we analyzed case-control cohort of 50 surgically resected HPV+ OPCs that later recurred (cases) and 50 tumors that were cured (controls) 1:1 matched for pathologic TNM stage, smoking history, and details of adjuvant therapy^31^. Matched features between cases and controls remained balanced between groups after exclusion of 3 samples containing HPV types not in the RNAScope probe set (Supplementary Table 3). The RNAscope-negative cell fraction within tumor regions was digitally quantified in the clinical OPC samples (Fig. 8a), which, in contrast to PDXs, included numerous nonmalignant cells in tumor areas comprised mostly of tumor infiltrating leukocytes (TILs). Thus, %TIL content in an adjacent section for each OPC was measured by CD45 immunohistochemistry (IHC) and used to estimate the HPV low OPC cell fraction by subtracting %TILs from % RNAscope-negative cells. The detailed analysis pipeline is illustrated in Fig. 8b. The Youden index cut-point for %HPV-low cells that optimally stratified recurrence status favorably discriminated between cases and controls based on area under the ROC curve (AUC)=0.784 (p<0.0001, OR=12.131, 95% CI=1.6-126.1) (Fig. 8c, top). The cases with locoregional failures (n=14) and their matched controls were also excluded to focus on tumors with only distant metastastic recurrence, which comprised most of the cases. Doing so further enhanced recurrence discrimination (AUC=0.808, p<0.0001, OR=28.283, 95% CI=21.4-688.8) (Fig. 8c, bottom). These differences were highly prognostic of overall survival stratified by the cut point for HPV-low cell fraction size that best discriminates recurrence risk (Fig 8d). We further evaluated any additional contributions of clinical and pathologic traits not matched between cases and controls to recurrence risk. Comparing these features (Supplementary Table 3) by univariate logistic regression identified lymphovascular space invasion (LVSI), median age, and the number of pathologically positive lymph nodes as statistically different between groups. Combining these features with HPV-low cell fraction by multivariate logistic regression (Table 2) showed that only HPV-low cells maintained significance (P < 0.001), supporting the score’s independent predictive utility. Together, these results show our findings linking HPV-low cell functions to cisplatin resistance to be generalizable to recurrence prediction and survival prognosis in HPV+ OPC patients receiving surgery followed by pathology-guided adjuvant therapy.

**Figure 8:**
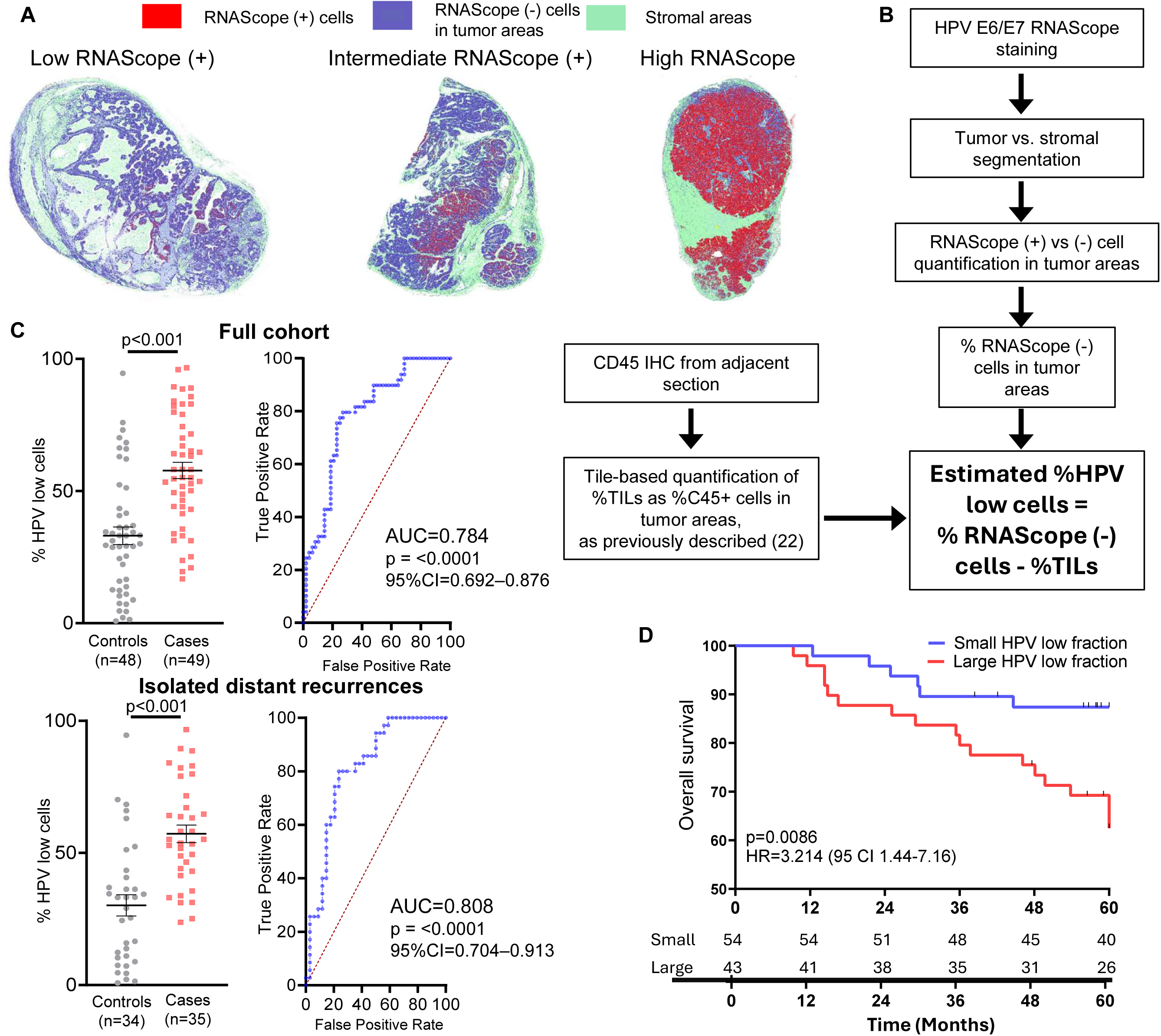
An expanded HPV low OPC cell fraction is associated with cancer recurrence risk and poor survival. **(A)** Representative segmentation of RNAscope staining micrographs (4x) for stromal regions and HPV E6/E7 RNAScope positive vs. negative cells in tumor regions using QuPath. **(B)** Algorithm for estimating HPV low OPC cell fraction size by combining RNAScope analysis with CD45 IHC-based quantification of %TILs in tumor areas of adjacent sections. **(C)** ROC curves distinguishing cases and controls based on HPV low cell fraction size, considering all recurrences (left) or the majority subset of recurrences comprised of isolated distant metastases (right). **(D).** Stratification of 5-year overall survival in the case-control cohort by the Youden index-based cut points for HPV low fraction size that optimally discriminate recurrence risk. P values are from log rank tests.

**Table 2:**
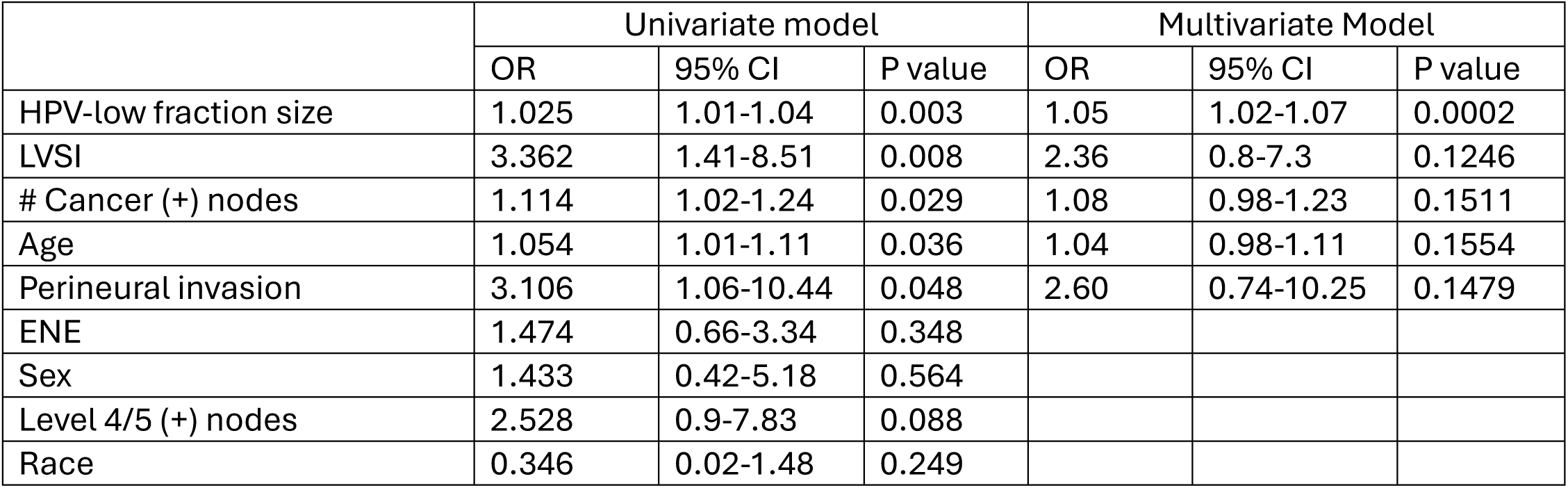
Univariate and multivariate logistic regression combining unmatched features of cases vs. controls with HPV-low cell fraction size for recurrence prediction. P-values and Odds ratio calculated by Logistic regression. CI = confidence interval; ENE = extranodal extension; LVSI = lymphovascular space invasion; OR = odds ratio

|  | Univariate model |  |  | Multivariate Model |  |  |
|---|---|---|---|---|---|---|
|  | OR | 95% CI | P value | OR | 95% CI | P value |
| HPV-low fraction size | 1.025 | 1.01-1.04 | 0.003 | 1.05 | 1.02-1.07 | 0.0002 |
| LVSI | 3.362 | 1.41-8.51 | 0.008 | 2.36 | 0.8-7.3 | 0.1246 |
| # Cancer (+) nodes | 1.114 | 1.02-1.24 | 0.029 | 1.08 | 0.98-1.23 | 0.1511 |
| Age | 1.054 | 1.01-1.11 | 0.036 | 1.04 | 0.98-1.11 | 0.1554 |
| Perineural invasion | 3.106 | 1.06-10.44 | 0.048 | 2.60 | 0.74-10.25 | 0.1479 |
| ENE | 1.474 | 0.66-3.34 | 0.348 |  |  |  |
| Sex | 1.433 | 0.42-5.18 | 0.564 |  |  |  |
| Level 4/5 (+) nodes | 2.528 | 0.9-7.83 | 0.088 |  |  |  |
| Race | 0.346 | 0.02-1.48 | 0.249 |  |  |  |

## DISCUSSION

This study offers novel molecular insights into an HPV-low tumor cell state in HPV+ malignancy. In addition, it uncovers a strong, independent association between HPV-low fraction size and OPC recurrence risk, providing a new basis for biomarker development. Unlike prior studies of comparable populations^5–8^, we present evidence that HPV-low cells retain stem-like transcriptional profiles and are not proliferatively impaired by senescence and/or terminal differentiation. We also show that the HPV-low OPC cells bioinformatically classified in most therapy-resistant PDXs are the most distinct transcriptionally, upregulating pathways that underlie adverse tumor biology, including RAS signaling, IL-6 signaling, and EMT. Such activation of alternate oncogenic signaling pathways in HPV-low cells could moderate overall addiction of HPV+ cancers to E6/E7-mediated signals. Pathways localizing to HPV-low cells were activated in signaling crosstalk predicted by CellChat, supporting cooperative roles for HPV-high and low fractions in therapy-refractory disease. Accordingly, the two fractions were maintained in stable proportions upon cisplatin exposure in therapy-resistant models. Our data also suggest that such crosstalk is sustained by certain genetic clones predisposed to enter the HPV-low state, supporting both genetic and epigenetic contributions to homeostasis of this population. Further delineating mechanisms of cooperativity and plasticity between HPV-low and high states may lead to identification of novel therapeutic strategies in treatment-refractory HPV+ malignancies.

Our findings linking HPV-low cells with therapy resistance, recurrence risk, and poor survival represent an advancement over prior work, which speculated on such roles from associations with reduced bulk level HPV mRNA. Specifically, lower bulk expression was linked to cisplatin resistance in subclonal derivatives of cancer cell lines and marginally worse survival in a small 28 patient cohort^7^. By contrast, we did not see a relationship between bulk level HPV mRNA and cisplatin responses of PDXs (Fig. 1d) or OPC recurrence risk in our case-control cohort^31^. In this present study, only the HPV-low state in the cisplatin-resistant subset of PDXs had markedly divergent host gene expression indicative of adverse cancer biology. Thus, bioinformatically processed scRNAseq definitions of HPV-low populations in treatment-sensitive models might be more artifactual, perhaps primarily representing rapid stochastic shifts in levels of low abundance, labile HPV mRNAs within single cells or random variability in RNAseq sensitivity near the threshold of detection. Consistent with this premise, a methodologically uniform, direct classification of the threshold for HPV E6/E7 detection in OPCs using a sensitive RNAScope assay was predictive of recurrence and prognostic of survival. Our validation of the association of HPV-low fraction size with cancer recurrence in a large, carefully curated case-control OPC cohort has a relatively direct pathway for clinical translation, given it is based on adaptation of a widely used RNAScope clinical assay.

Although our results do not prove HPV-low cells directly mediate recurrence, we identify multiple traits in this fraction with ability to promote an aggressive, recurrent phenotype. The HPV-low fraction not only retained stem-like states and tumor-propagating capacity but also contained fewer senescent cells than the HPV-high state. In this context, prior designations of comparable cells as “HPV off”^7^ may be misleading. The low HPV mRNA read counts seen in almost all OPC cells may easily fall below the threshold of detection by scRNAseq, causing a false impression of fully repressed HPV expression, which in experimental cancer models induces senescence and/or cell death^5^. By contrast, partial attenuation of E6/E7 expression below a detection threshold could mitigate DNA replication stress caused by the HPV E6/E7^32^, which can cause mitotic catastrophe, induce anti-tumor immunity, and augment genotoxic stress from standard therapy with radiation and cisplatin. Reduced E6/E7 could also limit oncogene-induced replicative senescence, explaining the larger senescent cell fraction shown here in the HPV-high population within PDXs and human OPCs. Hence, HPV low cells appear positioned to serve tumor progression in roles complementary to the higher biomass-generating capacity of HPV-high cells and to create a dynamic reservoir that repopulates the HPV-high cells driving rapid tumor growth.

Another feature setting this study apart is an improved ability to distinguish intrinsic HPV-low cell traits, which in prior work have been difficult to distinguish from immune microenvironment-dependent features. Recent spatial profiling of 7 human HPV+ OPCs showed HPV-low cells to be distributed between two compartments with distinct chemokine signaling. They were most enriched within hypoxic, neutrophil-rich CXCL8 foci but also appeared in CXCL9/10/11 foci with higher T cell content and prominent interferon signaling^5^. We show that HPV-low cells grown in NSG mice with no adaptive immunity and impaired innate immunity are highly similar to those in human OPCs and, show traits of those in both types of chemokine foci in aggregate. Specifically, their gene expression was indicative of both increased interferon signaling and tumor hypoxia. Both these features may help enforce the HPV-low state, given reported abilities of hypoxia ^33,34^ and interferon signaling ^35,36^ to repress HPV oncogene expression. The resulting decrease E6/E7 could subsequently enhance the proinflammatory milieu by derepressing cGAS/STING signaling^19–21^ in a feed-forward manner. Overall, our findings support HPV-low cells to be a core feature of therapy-resistant HPV+ malignancy whose existence is independent of adaptive immunity.

The immune-depleted milieu of PDXs enhanced inference of potential crosstalk among the OPC cells that might contribute to therapy resistance. The majority of predicted interactions in cisplatin-resistant PDXs were in cis among HPV-high cells. HPV-high cell-specific expression of tight junction components reflected their stronger epithelial identity relative to HPV-low cells, which upregulated EMT-related transcripts. However, a trans desmosomal interaction suggested HPV-high cells still tether to HPV-low cells expressing desmocollins, and loss of such attachment could also stimulate EGFR/c-Src signaling^29^ in HPV-high cells through desmogleins asymmetrically expressed on them. HPV-high cells also expressed secreted factor angiopoietin 4, which is involved in anoikis resistance, EMT, and metastasis by binding syndecans (Syndecan-1/4)^28^ and associated integrins found on both HPV-high and low cells. Syndecans activate pro-survival signals (Src, PI3K, MAPK) and upregulate MMP-1 to facilitate invasion^26–28^. Laminin interactions of integrins α2β1 and α6β1 on HPV-high cells predicted focal adhesion kinase (FAK) phosphorylation and activation of downstream signaling pathways (Src, PI3K)^37^, helping protect HPV-high cells from anoikis and promote survival. These interactions highlight contributions of HPV high/HPV low cell crosstalk to expansion of the HPV-low cell pool and related therapy resistance, underscoring utility of considering the distinct biology of each fraction in pursuing novel therapies.

## Methods

### PDX model growth, characterization, and classification of cisplatin responses

OPC PDX models driven by HPV16 were generated and passaged in the subcutaneous flanks of NSG mice, treated with cisplatin *in vivo,* and bulk mRNA sequenced as previously described^9,13,38^. NSG mice were used under Wistar Institute IACUC protocol #112652. Tumor volumes were measured as 0.5×length×width². Volume measurements at end of therapy were used to calculate T:C ratio (mean volume treated group/ mean volume control group). Cisplatin responses were evaluated using modified RECIST criteria, which were adapted to caliper-based tumor volume measurements in PDXs as previously described^39^. Treatment-naive OPC PDXs undergoing characterization by scRNAseq were harvested upon reaching 1cm^3^. Therapy responses were analyzed by scRNAseq after treatment of 0.5 cm^3^ PDXs with two doses of cisplatin three days apart or saline vehicle before harvesting at day 6.

### ScRNAseq: OPC cell preparation, mRNA sequencing, data filtration, and alignments

Freshly harvested xenograft tumors were trimmed of necrotic and adjacent mouse tissues and minced with surgical blades. Fragments were digested using the Tumor Dissociation Kit, human (Miltenyi Biotec) on a gentleMACS Dissociator for 30 min at room temperature. Digestion was stopped by adding an equal volume of medium containing 10% FBS, and the suspension was passed through 70-µm cell strainers and centrifuged. Debris was removed by density centrifugation with Debris Removal Solution per manufacturer’s instructions (Miltenyi Biotec). Dead and dying cells were then depleted using the Dead Cell Removal Kit (Miltenyi Biotec). To remove contaminating mouse stromal, endothelial, and hematopoietic cells, the viable fraction was subsequently processed with the Mouse Cell Depletion Kit (Miltenyi Biotec). Cell counts and viability were determined by trypan blue exclusion. Libraries were prepared with 10X Genomics Chromium Single cell 3’ (v3 chemistry) and sequenced on an Illumina NovaSeq 6000 platform for 10,000 UMIs per cell. A reference combining human and mouse genome (GRCm38) was downloaded from 10XGenomics to identify and exclude mouse cells. Cells with >75% UMIs aligned with mouse genome were identified as mouse origin and removed from further analysis. A custom reference genome combining the human genome (GRCh38) and HPV16 reference genome downloaded from PAVE^40^ was built using the Cell Ranger mkref protocol. Gene count matrices were generated for custom GRch38+HPV16 and demultiplexed using Cell Ranger. Low-quality OPC cells were excluded in Seurat v4.2 using cutoffs of nFeature <4,000 cap and percent.mt >25. Expression values across all samples were combined and normalized together, and an integrated object was created using Seurat the IntegrateLayers function. Dimension reduction and cell clustering were performed for the first 20 principal components. A nearest neighbor graph was constructed by calculating the neighborhood overlap using FindNeighbors. Unsupervised cell clustering was performed by Louvain algorithm in Find Clusters. To visualize the resulting clusters, UMAP dimension reduction using the same 20 principal components. HPV-aligned transcript counts were retained as cell-level metadata and excluded from the expression matrix used for downstream analyses. Raw human HPV+ OPC scRNAseq data from prior studies was downloaded from GEO accession numbers GSE182227 and GSE164690. Human samples were processed to retain high-quality cells using identical prefiltration criteria for nfeatures, percent.mt as PDX models. The samples annotated as HPV16 + were retained for further analysis. Cell types were determined using SingleR^41^ with HumanPrimaryCellAtlasData from CellDex^42^.

### Assignment of HPV low and HPV high OPC cell states in scRNAseq of PDXs and patient tumors

For each tumor, a range of ordinal cutoffs for total HPV transcript counts were tested. To avoid unstable estimates resulting from small group sizes, cutoffs resulting in either group containing fewer than 100 cells were excluded from analysis. Fractional read counts after data normalization were rounded to nearest ordinal values. OPC cells were divided into two groups using each possible cutoff, and the two groups were re-embedded by UMAP. Euclidean distances between centroids of each group in the UMAP space were computed, as detailed in Supplementary Methods. Cutoffs yielding maximal centroid distances were selected as optimal boundaries for separating HPV low from HPV high cells.

### Analysis of OPC cell gene expression, developmental states, subclones, and cell-cell interactions

The senescent OPC cell fraction was quantified by SenePy^16^ using the basal tongue epithelial senescence hub per the manual. Cell lineage commitment and differentiation states were evaluated using the classification pipeline in CytoTrace2^17^. Large-scale copy-number alterations were inferred with CopyKAT^23^ from raw UMI matrices after excluding mRNAs detected in fewer than five cells window size 50 and segmentation sensitivity of 0.15. CNAs were called in 1-Mb bins, clustered hierarchically (Euclidean distance, Ward’s linkage), and segmented by Bayesian inference with a 25-gene smoothing window. Differential gene expression between HPV low and HPV high cells were assessed using MAST in Seurat^43^. Genes were ranked by average log_2 f_old-change and ranked lists tested against the MSigDB Hallmark gene sets^15^ (n=50) by gene set enrichment analysis (GSEA) using AUCCell^44^. Pseudobulk profiles were generated using normalized counts by the Seurat PseudobulkExpression function, treating each biological sample as an independent replicate for HPV-low and HPV-high cells individually. Interactions between HPV-high and HPV-low cells were inferred by CellChat^24^ using normalized expression matrices. Communication probabilities and p-values are inferred using triMean method with 10,000 bootstraps for individual sample. Interactions with p-values <0.2 were aggregated to compare interaction numbers and directionality between HPV-low and HPV-high cells. Pathway-level networks and central signaling roles were examined to identify communication profiles.

### Analysis of HPV+ OPC tissue samples in the case-control cohort

We have previously detailed a single institution cohort of 50 surgically resected p16+ OPC cases that later recurred plus 1:1 matched control tumor that had follow up beyond the latest recurrence event. Briefly, cases and controls were matched based on 8^th^ edition AJCC TNM staging, smoking history, and details of adjuvant therapy^31^. Bulk RNA sequencing of pre-treatment tumors from these patients is described previously^31^. The same FFPE tumor sections previously used for bulk RNA sequencing were probed by RNAScope for HPV E6/E7 from four high-risk HPV genotypes (16, 18, 31, 33) using a standard clinical assay for confirmation of HPV relatedness, as detailed previously^45^. A small number of case and control tumors proven in this prior study to have false negative RNAScope results due to the probe set lacking their high-risk HPV genotype (n=3) were excluded from analysis. RNAScope-stained slides were digitized on a 3DHISTECH Panoramic scanner and analyzed using QuPath (v0.5.1)^46^. Total tissue area was segmented using a pixel intensity threshold, and an object classifier was trained individually on each sample to distinguish tumor parenchyma from stroma. Cells in tumor areas were segmented using default parameters, and individual cells were classified as HPV positive if the maximum 3,3’-diaminobenzidine optical density within the cell exceeded 0.1 %RNAScope-negative cells were calculated as 100-%HPV positive cells. Non-malignant cells in tumor areas were almost all tumor-infiltrating leukocytes (TILs), which were identified by CD45 IHC in the adjacent FFPE section and quantified using QuPath as previously detailed^31^. Therefore, the HPV low tumor cell fraction was approximated by subtracting %TILs from %RNAScope-negative cells.

### Statistics

Analyses were performed using Rv4.2 and GraphPad Prism. Receiver operating characteristic (ROC) analyses were performed using pROC. P-values for area under the ROC curve (AUC) were calculated from the z ratio using the normal distribution. Survival analyses were performed in R using Survminer package. All tests were two-tailed, and statistical significance was defined as p<0.05 unless otherwise noted. Statistical tests used for each comparison are specified in figure legends.

## Acknowledgements

D.B, P.G., and J.B.J. were supported by NIH R01DE034056, UH2CA267502, and R01DE027185. The funders had no role in the study design, data collection and analysis, decision to publish or preparation of this manuscript. Work was also supported by the Alpha Epsilon Fund, the Breakthrough Challenge Foundation, the Stephen and Susan Kelly Family Fund for Head and Neck Cancers, and the William and Greta Lydecker Fund.

## Author Contributions

Conceptualization (L.R., M.K.S, D.B., J.G., A.D., N.A., S.M.S., I.M.M.); Formal analysis (L.R, M.R.P., M.K.S, J.B.J, J.G., P.G.); Funding acquisition (D.B.); Investigation (L.R, M.R.P, M.K.S, D.B., J.G., P.R., R.M.B., J.B.J., N.A.); Project administration (D.B.); Supervision (M.K.S., D.B.); Writing-original draft (L.R., M.R.P., M.K.S., D.B.); Writing-review and editing (all authors)

## Human Ethics and Consent to Participants declaration

Tissues used for PDXs were collected from patients undergoing primary surgical resections at the Hospital of the University of Pennsylvania (protocol #417200) or the Philadelphia VA Medical Center (protocol #01090) as previously detailed^38^. HPV+ OPCs used for RNA scope are from a previously detailed single institution cohort (protocol number #833890)^31^.

## Data availability declaration

ScRNA sequencing data generated in this study is in the process of being submitted and the accession number is pending. Bulk RNA sequencing from the PDXs used in this study is available through GEO database under the accession number GSE193388^13^. RNAseq for the case control cohort was generated in a prior study ^47^ and is deposited in dbGAP Study ID phs003889, title: Pursuing molecular biomarkers to guide adjuvant therapy for HPV head and neck cancers after transoral robotic surgery. ScRNAseq from human HPV+ OPCs was generated in two prior studies^7,8^ and is available in through GEO database under the accession numbers GSE182227 and GSE164690.

