## Supplementary figures and images for "An expanded, phenotypically distinct tumor cell subset with low HPV mRNA increases recurrence risk"

### Supplementary Figure 1

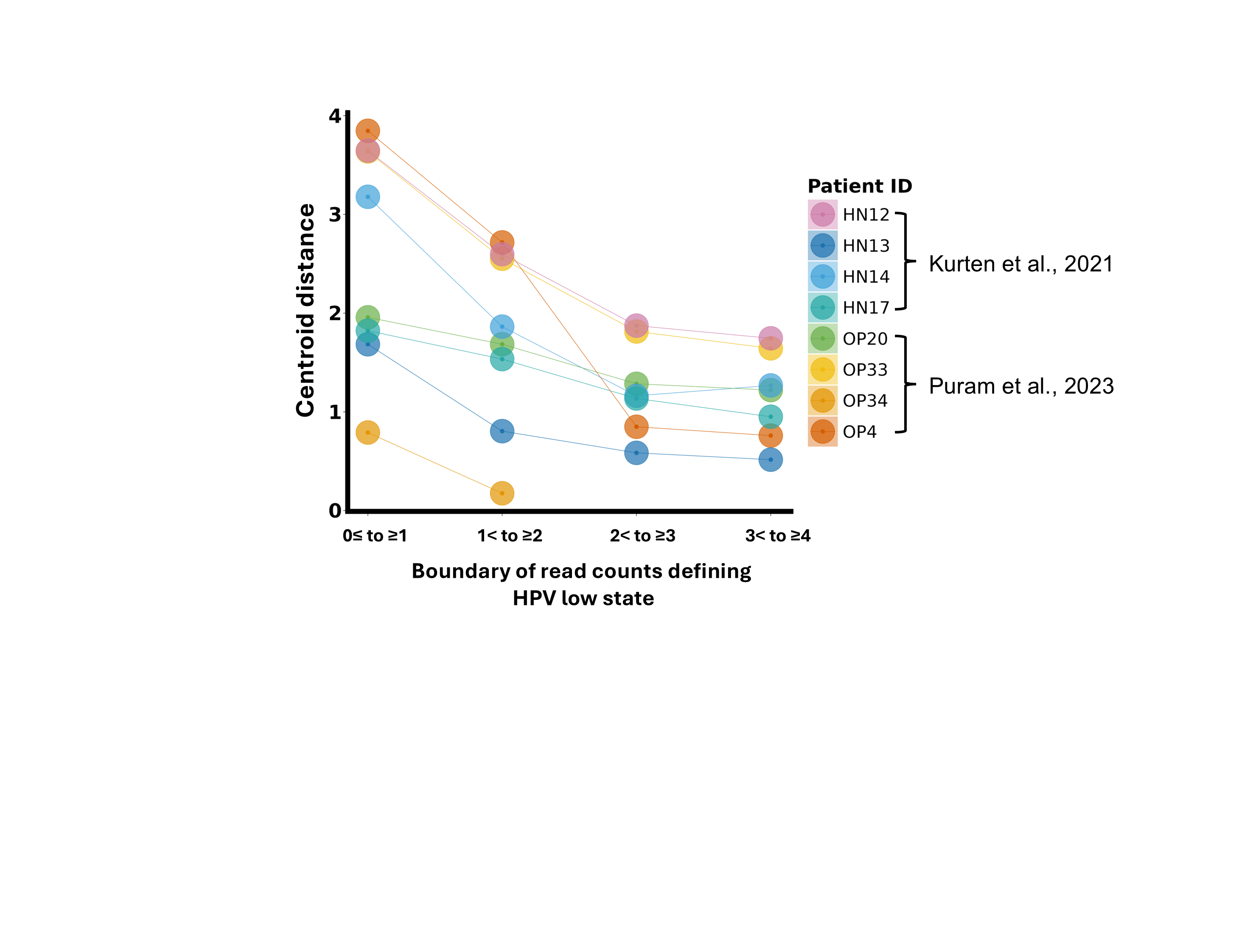

### Supplementary Figure 2

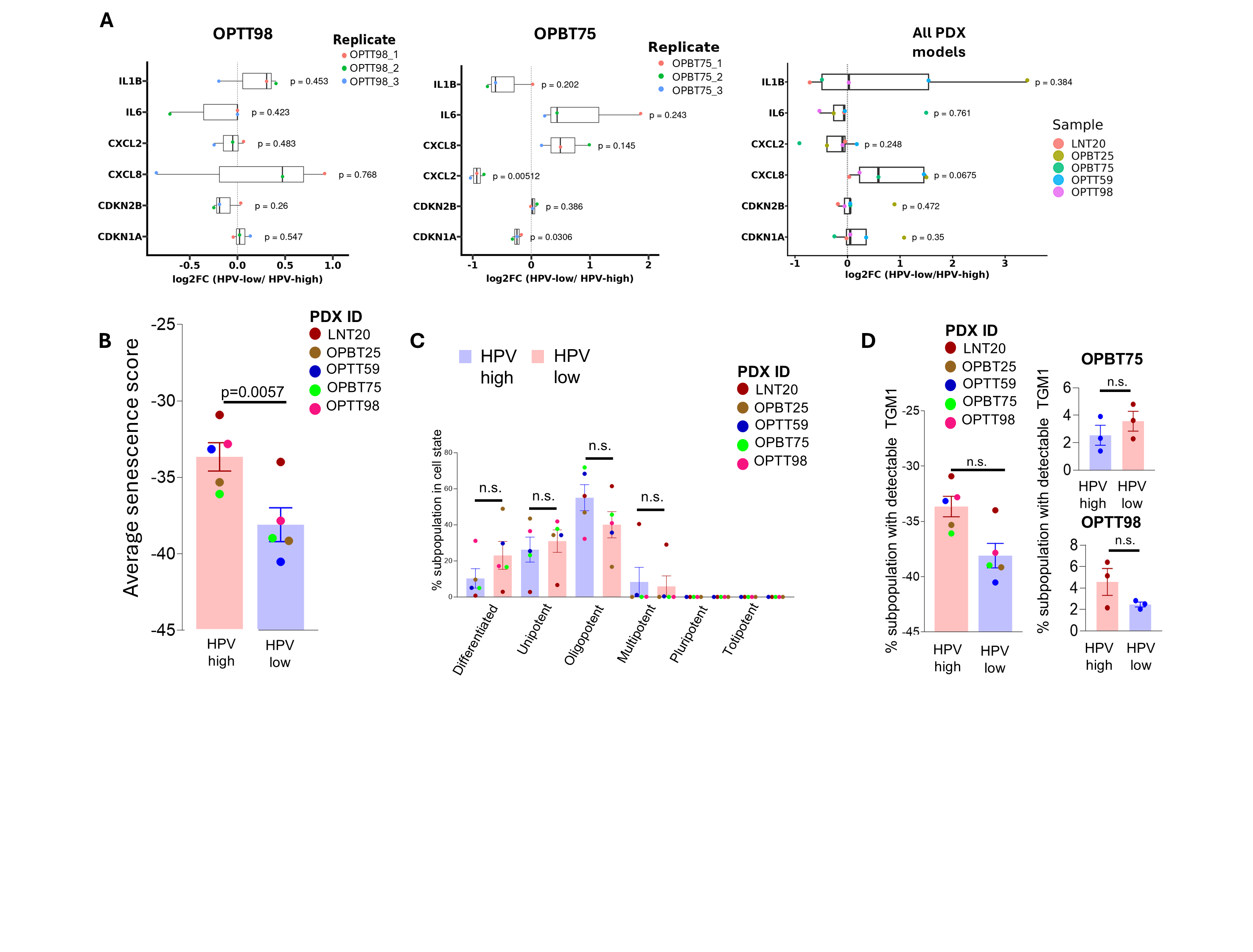

### Supplementary Figure 3

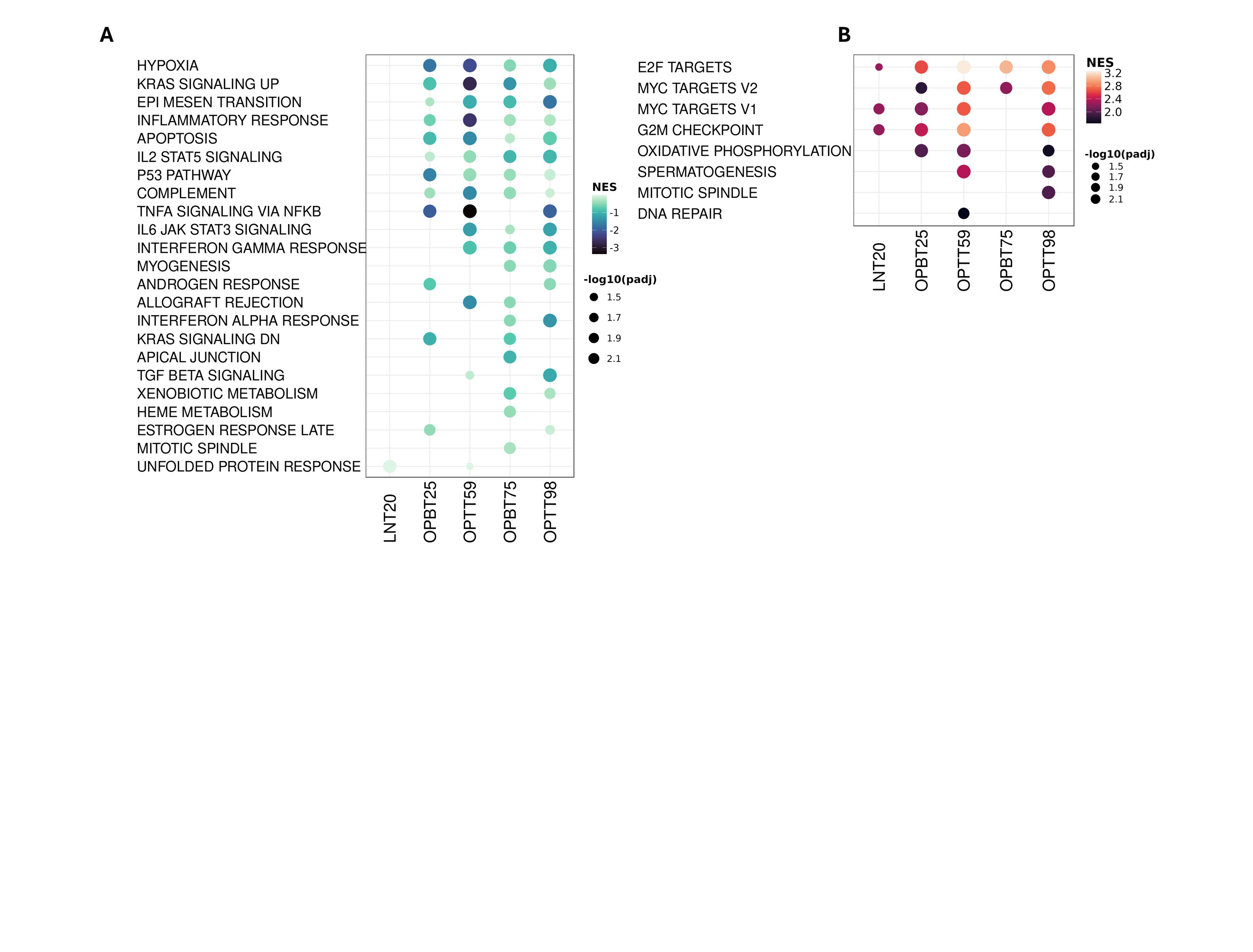

### Supplementary Figure 4

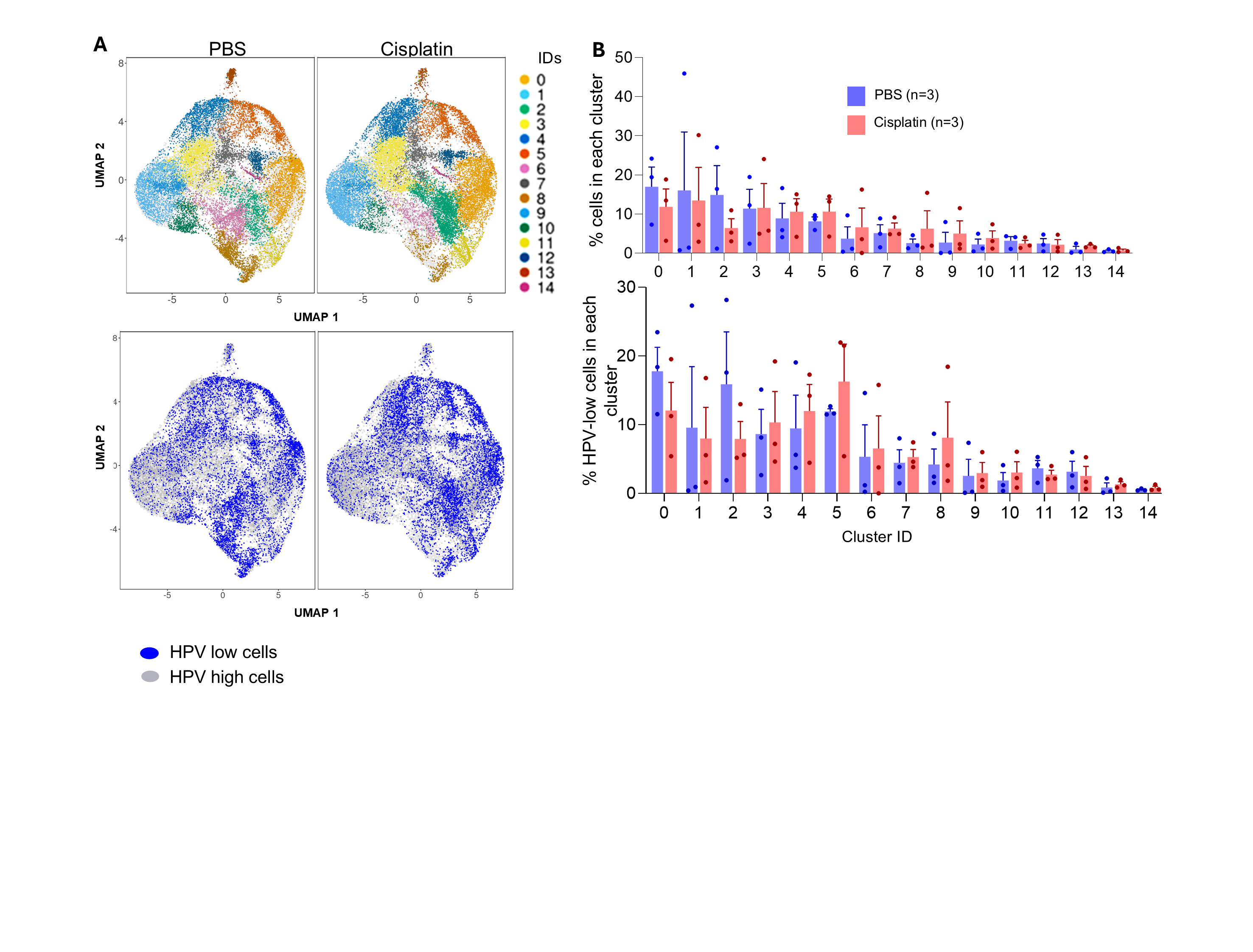

### Supplementary Figure 5

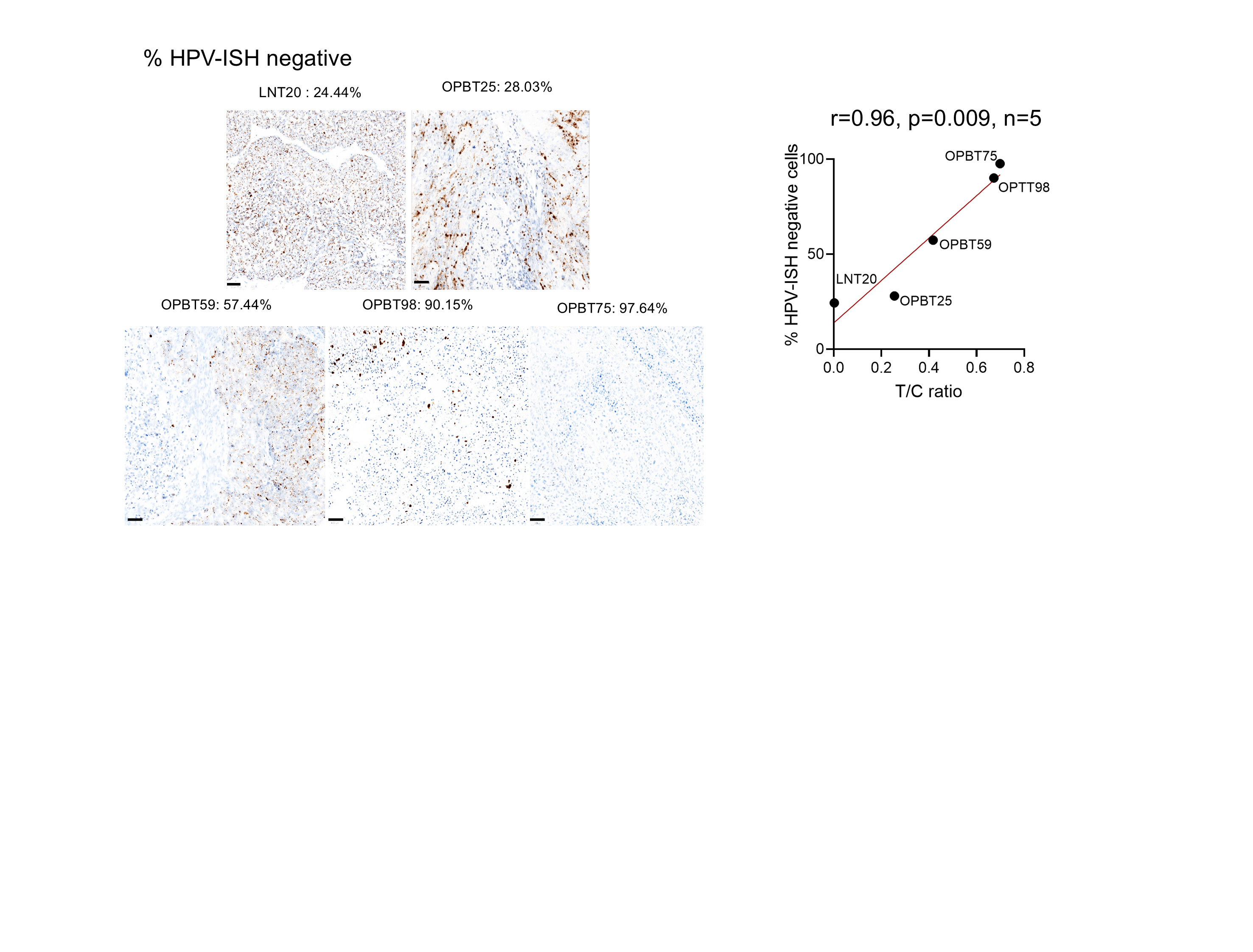
