## Supplementary information for "An expanded, phenotypically distinct tumor cell subset with low HPV mRNA increases recurrence risk"

**Supplementary Figure 1**: Distribution of centroid distances for evaluable, publicly available HPV+ OPC samples with >1,000 cells.

**Supplementary Figure 2**: **Senescence and stem-like features in HPV low and high cells. (A)** Relative SASP and CDK inhibitor expression as $\log_2$ foldchange (HPV low / HPV high) of pseudobulk levels in PDX models. **(B)** SenePy-derived senescence score in HPV-high versus HPV-low cells across PDX models. **(C)** Proportion of HPV high (blue bar) and HPV low (red bar) cells distributed across CytoTRACE2-defined developmental states in PDXs. Bars represent mean ± SEM. **(D)** Proportion of HPV high and HPV low cells expressing TGM1 across all five PDX models (left) and individually for OPBT75 and OPTT98 with biological replicates (right). Significance was assessed using paired t-tests throughout.

**Supplementary Figure 3**: **Pathways significantly downregulated (left) and upregulated (right) in at least 1 PDX.** GSEA compares pseudobulk aggregates of HPV-low **(A)** to HPV-high **(B)** populations using the Hallmark and KEGG pathways. Dot sizes reflect adjusted P values, and color intensity shows normalized enrichment score (NES). Differential pathway expression is designated based on adjusted $p < 0.05$ by the Benjamini–Hochberg method.

**Supplementary Figure 4**: **Stability of HPV-low and HPV-high cells upon treating OPTT98.** (A) UMAP clustering of treated and control OPTT98 OPC cells (top) overlaid with HPV-low and high cell identities (bottom). (B) Proportions of total OPC cells (top) and HPV-low cells (bottom) in each UMAP cluster in control vs. treated OPTT98 PDXs, with lack of significant differences observed using Sidak's multiple comparisons test.

**Supplementary Figure 5: Percentage of E6/E7 RNAScope (-) OPC cells in PDXs correlates with cisplatin resistance.** Micrographs on the left are representative RNAScope images of the 5 PDX tumors harvested at comparable sizes early in their growth curves, with percent of RNAScope (-) cells in tumor areas indicated for each. The Pearson correlation analysis on right shows relationship of these percentages to T:C ratios for cisplatin. Images are 20x, Bars=50 μm.

**Supplementary table 1:** Complete list of interactions inferred by CellChat between HPV high and HPV low cells in either OPBT75 or OPTT98. P-adjusted values calculated by Benjamini-Hochberg (BH) method incorporated in fGSEA. NES= Normalized enrichment score

| PDX | Apoptosis | | Apoptosis-p53 | |
|---|---|---|---|---|
| | NES | p adjusted value | NES | p adjusted value |
| LNT20 | 1.084417 | 0.504 | 1.084417 | 0.504 |
| OPBT25 | 1.881869 | 0.021 | 1.406407 | 0.053 |
| OPBT75 | 1.496489 | 0.041 | 1.290516 | 0.202 |
| OPTT59 | 2.223664 | 0.008 | 1.382756 | 0.050 |
| OPTT98 | 1.734425 | 0.015 | 1.484899 | 0.018 |

**Supplementary Table 2**: Features matched between cases and controls. P-values are inferred using triMean method incorporated in CellChat with 10,000 bootstraps

**Supplementary Table 3:** Features not matched between cases and controls. Categorical p-values calculated by Pearson's chi-square or Fisher exact test. Continuous p-values calculated by Wilcox-Rank sum test with continuity correction. CRT = chemoradiation; RT = radiation therapy; LVSI = lymphovasciular space invasion; ENE = extranodal extension; BMI = body mass index

**Supplementary methods: Centroid-distance optimization of HPV-low/HPV-high thresholds**

The centroid (C) is the mean coordinate vector for all cells categorized to the HPV-low or HPV-high group based on the viral read cutoff values within the UMAP space

$$C_{HPV-low} = \frac{1}{n_{HPV-low}} \sum_{i=1}^{n_{HPV-low}} x_{i,d}$$

$$C_{HPV-high} = \frac{1}{n_{HPV-high}} \sum_{j=1}^{n_{HPV-high}} y_{j,d}$$

The centroid distance (D) for individual viral expression cutoffs is defined for N dimensions

$$D(C_{HPV-low}, D_{HPV-high}) = \sqrt[2]{\sum_{d=1}^{N} (C_{HPV-low,d} - C_{HPV-high,d})^{2}}$$

Where:

- $C_{HPV\text{-}low}$ and $C_{HPV\text{-}high}$ are the centroids for HPV-low and HPV-high populations respectively
- $n_{HPV\text{-}low}$ and $n_{HPV\text{-}high}$ are total number of cells in HPV-low and HPV-high cell clusters respectively
- $x_i$ is the coordinate vector of cell *i* in HPV-low cell cluster
- $y_j$ is the coordinate vector of cell *j* in HPV-high cell cluster
- D is the centroid distance between cell populations defined as HPV-low and HPV-high
- d is the index representing an individual dimension
- N is the number of principle component dimensions identified
