## Supplementary table 3 for "An expanded, phenotypically distinct tumor cell subset with low HPV mRNA increases recurrence risk"

| Source | Target | Ligand | Receptor | P value | Signaling pathway | Type of interaction |
|---|---|---|---|---|---|---|
| **Cis interactions** | | | | | | |
| **OPBT75** | | | | | | |
| High | High | OCLN | OCLN | 0.000789 | OCLN | Cell-Cell Contact |
| High | High | 12oxoLTB4 | LTB4R | 0.00081 | 12oxoLTB4 | Non-protein Signaling |
| High | High | CDH3 | CDH3 | 0.000877 | CDH | Cell-Cell Contact |
| High | High | MPZL1 | MPZL1 | 0.000969 | MPZ | Cell-Cell Contact |
| High | High | GJA1 | GJA1 | 0.001101 | GAP | Cell-Cell Contact |
| High | High | AREG | EGFR | 0.001304 | EGF | Secreted Signaling |
| High | High | CDH1 | CDH1 | 0.001987 | CDH | Cell-Cell Contact |
| High | High | PLAU | PLAUR | 0.002046 | PLAU | Secreted Signaling |
| High | High | HBEGF | EGFR | 0.00226 | EGF | Secreted Signaling |
| High | High | ANGPTL4 | SDC4 | 0.002962 | ANGPTL | Secreted Signaling |
| High | High | CDH1 | ITGA2_ITGB1 | 0.003626 | CDH1 | Cell-Cell Contact |
| High | High | GRN | SORT1 | 0.00365 | GRN | Secreted Signaling |
| High | High | ANGPTL4 | SDC1 | 0.005816 | ANGPTL | Secreted Signaling |
| High | High | DSC2 | DSG2 | 0.008869 | DESMOSOME | Cell-Cell Contact |
| High | High | LAMB3 | ITGA3_ITGB1 | 0.012789 | LAMININ | ECM-Receptor |
| High | High | DSC3 | DSG2 | 0.01289 | DESMOSOME | Cell-Cell Contact |
| High | High | LAMB3 | ITGA2_ITGB1 | 0.013555 | LAMININ | ECM-Receptor |
| High | High | CLDN1 | CLDN1 | 0.013695 | CLDN | Cell-Cell Contact |
| High | High | CD99 | CD99 | 0.022185 | CD99 | Cell-Cell Contact |
| High | High | LAMB3 | CD44 | 0.025108 | LAMININ | ECM-Receptor |
| High | High | LAMB3 | ITGA6_ITGB1 | 0.026384 | LAMININ | ECM-Receptor |
| High | High | LAMB3 | ITGA6_ITGB4 | 0.033484 | LAMININ | ECM-Receptor |
| High | High | DSC3 | DSG3 | 0.04657 | DESMOSOME | Cell-Cell Contact |
| High | High | F11R | F11R | 0.048568 | JAM | Cell-Cell Contact |
| High | High | PPIA | BSG | 0.167146 | CypA | Secreted Signaling |
| **OPTT98** | | | | | | |
| High | High | CLDN1 | CLDN1 | 0.001487 | CLDN | Cell-Cell Contact |
| High | High | EFNA1 | EPHA2 | 0.001865 | EPHA | Cell-Cell Contact |
| High | High | OCLN | OCLN | 0.002558 | OCLN | Cell-Cell Contact |
| High | High | LAMC2 | CD44 | 0.003153 | LAMININ | ECM-Receptor |
| High | High | ANGPTL4 | SDC4 | 0.00639 | ANGPTL | Secreted Signaling |
| High | High | ANGPTL4 | SDC1 | 0.012656 | ANGPTL | Secreted Signaling |
| High | High | LAMC2 | ITGA2_ITGB1 | 0.013722 | LAMININ | ECM-Receptor |
| High | High | LAMB3 | CD44 | 0.013878 | LAMININ | ECM-Receptor |
| High | High | LAMC2 | ITGA6_ITGB4 | 0.014785 | LAMININ | ECM-Receptor |
| High | High | LAMC2 | ITGA6_ITGB1 | 0.018704 | LAMININ | ECM-Receptor |
| High | High | CD99 | CD99 | 0.044797 | CD99 | Cell-Cell Contact |
| High | High | LAMB3 | ITGA2_ITGB1 | 0.058297 | LAMININ | ECM-Receptor |
| High | High | LAMB3 | ITGA6_ITGB4 | 0.062593 | LAMININ | ECM-Receptor |
| High | High | LAMB3 | ITGA6_ITGB1 | 0.07818 | LAMININ | ECM-Receptor |
| Low | Low | JAG1 | NOTCH3 | 0.001275 | NOTCH | Cell-Cell Contact |
| Low | Low | PTN | SDC1 | 0.009999 | PTN | Secreted Signaling |
| Low | Low | PTN | NCL | 0.012707 | PTN | Secreted Signaling |
| Low | Low | LAMB3 | CD44 | 0.013832 | LAMININ | ECM-Receptor |

| | | | | | | |
|---|---|---|---|---|---|---|
| Low | Low | DSC3 | DSG2 | 0.022729 | DESMOSOME | Cell-Cell Contact |
| Low | Low | DSC3 | DSG1 | 0.02507 | DESMOSOME | Cell-Cell Contact |
| Low | Low | DSC2 | DSG1 | 0.02949 | DESMOSOME | Cell-Cell Contact |
| Low | Low | DSC3 | DSG3 | 0.147333 | DESMOSOME | Cell-Cell Contact |
| Low | Low | DSC2 | DSG3 | 0.16956 | DESMOSOME | Cell-Cell Contact |
| **Trans interactions** | | | | | | |
| **OPBT75** | | | | | | |
| Low | High | CLDN1 | CLDN1 | 0.005802 | CLDN | Cell-Cell Contact |
| Low | High | DSC2 | DSG2 | 0.009175 | DESMOSOME | Cell-Cell Contact |
| Low | High | LAMB3 | ITGA3_ITGB1 | 0.012862 | LAMININ | ECM-Receptor |
| Low | High | LAMB3 | ITGA2_ITGB1 | 0.013633 | LAMININ | ECM-Receptor |
| Low | High | LAMB3 | CD44 | 0.02525 | LAMININ | ECM-Receptor |
| Low | High | LAMB3 | ITGA6_ITGB1 | 0.026533 | LAMININ | ECM-Receptor |
| Low | High | DSC2 | DSG3 | 0.033477 | DESMOSOME | Cell-Cell Contact |
| Low | High | LAMB3 | ITGA6_ITGB4 | 0.033672 | LAMININ | ECM-Receptor |
| Low | High | PPIA | BSG | 0.16002 | CypA | Secreted Signaling |
| High | Low | ANGPTL4 | SDC4 | 0.001488 | ANGPTL | Secreted Signaling |
| High | Low | ANGPTL4 | SDC1 | 0.004671 | ANGPTL | Secreted Signaling |
| High | Low | CLDN1 | CLDN1 | 0.005802 | CLDN | Cell-Cell Contact |
| **OPTT98** | | | | | | |
| Low | High | Cholestero | RORA | 0.001142 | Cholesterol | Non-protein Signaling |
| Low | High | JAG1 | NOTCH3 | 0.001397 | NOTCH | Cell-Cell Contact |
| Low | High | EFNA1 | EPHA2 | 0.001586 | EPHA | Cell-Cell Contact |
| Low | High | PTN | SDC4 | 0.005174 | PTN | Secreted Signaling |
| Low | High | LAMC2 | ITGA2_ITGB1 | 0.006954 | LAMININ | ECM-Receptor |
| Low | High | PTN | SDC1 | 0.01026 | PTN | Secreted Signaling |
| Low | High | PTN | NCL | 0.011462 | PTN | Secreted Signaling |
| Low | High | DSC3 | DSG2 | 0.024335 | DESMOSOME | Cell-Cell Contact |
| Low | High | DSC2 | DSG2 | 0.028629 | DESMOSOME | Cell-Cell Contact |
| Low | High | LAMB3 | ITGA2_ITGB1 | 0.046528 | LAMININ | ECM-Receptor |
| Low | High | LAMB3 | ITGA6_ITGB1 | 0.062665 | LAMININ | ECM-Receptor |
| Low | High | DSC3 | DSG3 | 0.13724 | DESMOSOME | Cell-Cell Contact |
| High | Low | LAMC2 | CD44 | 0.003983 | LAMININ | ECM-Receptor |
| High | Low | CD46 | JAG1 | 0.005026 | CD46 | Cell-Cell Contact |
| High | Low | LAMC2 | ITGA2_ITGB1 | 0.007998 | LAMININ | ECM-Receptor |
| High | Low | ANGPTL4 | SDC1 | 0.012335 | ANGPTL | Secreted Signaling |
| High | Low | LAMC2 | ITGA6_ITGB4 | 0.014539 | LAMININ | ECM-Receptor |
| High | Low | LAMC2 | ITGA6_ITGB1 | 0.017157 | LAMININ | ECM-Receptor |
| High | Low | LAMB3 | CD44 | 0.017482 | LAMININ | ECM-Receptor |
| High | Low | DSC3 | DSG1 | 0.021346 | DESMOSOME | Cell-Cell Contact |
| High | Low | DSC2 | DSG1 | 0.026375 | DESMOSOME | Cell-Cell Contact |
| High | Low | LAMB3 | ITGA2_ITGB1 | 0.034632 | LAMININ | ECM-Receptor |
| High | Low | LAMB3 | ITGA6_ITGB4 | 0.061603 | LAMININ | ECM-Receptor |
| High | Low | LAMB3 | ITGA6_ITGB1 | 0.072076 | LAMININ | ECM-Receptor |
