## Supplementary table 3 for "An expanded, phenotypically distinct tumor cell subset with low HPV mRNA increases recurrence risk"

Overall cohort

| | Variable | Categories | R (n=49) | C (n=48) | P-value |
|---|---|---|---|---|---|
| Matched features | Overall pathologic stage | Stage I | 25 | 24 | 1.000 |
| | | Stage II | 24 | 24 | |
| | Pathologic T-stage | pT1 | 14 | 11 | 0.749 |
| | | pT2 | 28 | 31 | |
| | | pT3 | 7 | 6 | |
| | Pathologic N-stage | pN0 | 5 | 5 | 0.954 |
| | | pN1 | 27 | 25 | |
| | | pN2 | 17 | 18 | |
| | Treatment | Surgery | 4 | 4 | 0.995 |
| | | Surgery + RT | 19 | 19 | |
| | | Surgery + CRT | 26 | 25 | |
| | Smoking history | ≤10 Pack/yr | 34 | 33 | 0.617 |
| | | >10 Pack/yr | 15 | 15 | |
| Unmatched features | Overall clinical stage | Stage I | 25 | 25 | 1.000 |
| | | Stage II | 24 | 23 | |
| | Clinical T-stage | cTx | 4 | 11 | 0.195 |
| | | cT1 | 12 | 8 | |
| | | cT2 | 26 | 26 | |
| | | cT3 | 3 | 2 | |
| | | cT4 | 4 | 1 | |
| | Clinical N-stage | cN0 | 5 | 5 | 0.939 |
| | | cN1 | 26 | 27 | |
| | | cN2 | 18 | 16 | |
| | Primary tumor site | Tonsil | 24 | 24 | 0.562 |
| | | Tongue base | 20 | 17 | |
| | | Overlap | 4 | 7 | |
| | | Unknown primary | 1 | | |
| | Level 4/5 (+) nodes | No | 36 | 42 | 0.138 |
| | | Yes | 13 | 6 | |
| | Margin status | Negative | 30 | 29 | 1.000 |
| | | Positive | 19 | 19 | |
| | LVSI | No | 26 | 38 | 0.013 |
| | | Yes | 23 | 10 | |
| | Perineural invasion | No | 36 | 43 | 0.075 |
| | | Yes | 13 | 5 | |
| | ENE | No | 26 | 26 | 1.000 |
| | | Yes | 23 | 23 | |
| | # Cancer (+) nodes | Mean | 5.735 | 3.229 | 0.403 |
| | | Range | 0-25 | 0-10 | |
| | Age | Median | 62 | 58 | 0.029 |
| | | Range | 32-84 | 34-69 | |
| | BMI | Median | 28.345 | 28.45 | 0.801 |
| | | Range | 19.49-37.3[illegible] | 20.48-58.32 | |
| | Sex | Male | 42 | 43 | 0.787 |
| | | Female | 7 | 5 | |

| | | | | | |
|---|---|---|---|---|---|
| | Race | White | 48 | 45 | 0.595 |
| | | Non-white | 1 | 3 | |
